# *Akkermansia muciniphila*-derived LPS links gut dysbiosis to pathogenic miR-21 signaling in experimental autoimmune encephalomyelitis

**DOI:** 10.64898/2026.07.06.736880

**Authors:** Manu Mallahalli, Hirohiko Hohjoh, Daiki Takewaki, Kimitoshi Kimura, Shinji Oki, Hiroshi Mori, Koji Hosomi, Jun Kunisawa, Atsushi Toyoda, Wakiro Sato, Takashi Yamamura

## Abstract

Multiple sclerosis (MS) is a chronic T cell-mediated autoimmune disease characterized by blood-brain barrier (BBB) disruption, neuroinflammation, and demyelination of the central nervous system (CNS). Emerging evidence links gut microbiota to disease pathogenesis, but the microbial factors that regulate pathogenic microRNA (miRNA) programs are largely unknown. Here, using experimental autoimmune encephalomyelitis (EAE, a MS mouse model), we investigated whether gut microbiota exacerbate EAE pathogenesis by modulating host miRNA expression. Antibiotic-induced depletion of the gut microbiota markedly attenuated EAE scores and reduced circulating inflammatory miRNAs, with miR-21 emerging as the dominant pathogenic candidate. Functional inhibition of miR-21 significantly ameliorated disease severity and reduced CNS T-cell infiltration. Mechanistically, miR-21 enhanced IL-17 and GM-CSF production by CD4⁺ T cells and promoted immune-cell entry into the CNS through endothelial activation and blood-brain barrier dysfunction. We identified a transient expansion of *Akkermansia muciniphila* during the prodromal phase of EAE that positively correlated with circulating miR-21 levels. Colonization of antibiotic-treated mice with *A. muciniphila* exacerbated EAE and increased serum miR-21, whereas monocolonization of germ-free mice was insufficient to induce systemic miR-21, indicating a requirement for an inflammatory host environment. Further analyses revealed that atypical lipopolysaccharides (LPS) derived from *A. muciniphila* induce epithelial miR-21 production through coordinated TLR2/TLR4 signaling. Circulating miR-21 subsequently promoted endothelial dysfunction through the TIMP3-ADAM17 pathway, facilitating pathogenic T-cell migration into the CNS. Importantly, circulating miR-21 was also elevated in patients with MS. Collectively, these findings identify a previously unrecognized *A. muciniphila*–LPS–miR-21 axis linking gut dysbiosis to neuroinflammation and suggest that host-derived miRNAs function as systemic mediators through which microbial signals influence CNS autoimmunity.

## Introduction

MicroRNAs (miRNAs) are small, non-coding RNAs that regulate gene expression post-transcriptionally by either activating or silencing target genes [1, 2]. Increasing evidence highlights their roles in disease pathogenesis and their potential as diagnostic biomarkers. In multiple sclerosis (MS), a demyelinating autoimmune disease, autoreactive immune cells are activated in the periphery and then infiltrate the central nervous system (CNS) through the Blood-Brain Barrier (BBB). Proinflammatory T helper 17 cells are the principal mediators of this CNS inflammation, yet the precise mechanisms that drive the initial differentiation of naive T cells into autoreactive lineages remain highly complex [3]. Because several conserved miRNAs govern MS-related immune cascades, dysregulated serum miRNA profiles are frequently observed in MS patient cohorts [4–7]. Among these, microRNA-21a-5p (miR-21) has attracted particular interest because of its established role in inflammatory responses and autoimmune disease. However, despite extensive characterization of disease-associated miRNAs, the mechanisms governing their production and the origin of circulating pathogenic miRNAs remain poorly understood [8–11]. Identifying upstream factors that regulate pathogenic miRNA expression is, therefore, a critical step toward understanding MS pathogenesis.

Accumulating evidence, including our own previous work, demonstrates that the composition of the gut microbiota in patients with MS is significantly altered compared to healthy controls [12–15]. Gut inflammation and dysbiosis are increasingly recognized features of both MS and experimental autoimmune encephalomyelitis (EAE), a murine model classically induced using myelin oligodendrocyte glycoprotein (MOG) peptides. In EAE, the MOG-induced T cell acts as the primary cellular engine of autoimmunity, while the breach of the blood-brain barrier (BBB) serves as the critical gatekeeping event that transitions systemic immune activation into CNS tissue destruction[16–18]. Notably, germ-free or antibiotic-treated mice exhibit an impaired development of inflammatory T helper cells, rendering them highly resistant to EAE relative to specific-pathogen-free (SPF) counterparts [19, 20]; however, recolonization of these mice with SPF microbiota fully restores disease susceptibility [21]. Furthermore, in adoptive transfer models, myelin-specific TCRMOG T cells have been shown to transiently infiltrate the gut prior to CNS migration, a process that alters local microbial composition and enhances the pathogenicity of the infiltrating cells [22]. Collectively, these findings establish the gut microbiota as a critical regulator of autoimmune neuroinflammation. Nevertheless, whether specific microbial signals drive this CNS pathology by modulating host microRNA (miRNA) production remains to be elucidated. Emerging evidence indicates that gut microbes can regulate host miRNA expression through direct interactions with epithelial and immune cells. Microbial-associated molecular patterns, including lipopolysaccharides (LPS), activate Toll-like receptor (TLR)-dependent signaling pathways that influence transcriptional programs controlling miRNA expression [23]. Several studies have shown that microbial colonization and bacterial products alter the abundance of inflammation-associated miRNAs in intestinal epithelial cells, myeloid cells, and circulating extracellular vesicles, suggesting that disease-associated gut microbes may promote autoimmune pathology through miRNA-dependent mechanisms [24, 25].

Among gut microbes associated with MS, *Akkermansia muciniphila* has emerged as one of the most consistently reported species [26–29]. We and others have reported elevated *A. muciniphila* levels in patients with relapsing-remitting MS (RRMS). However, its immunological role remains unresolved. Some studies report anti-inflammatory effects through regulatory T-cell induction, whereas others demonstrate that *A. muciniphila* isolated from MS patients promotes Th1-type responses in healthy peripheral blood mononuclear cells [27]. Similarly, different *A. muciniphila* strains exert distinct effects on inflammatory responses in EAE models [12, 30]. These observations suggest that *A. muciniphila* may influence host immunity through mechanisms that remain poorly defined.

To address this question, we investigated whether an MS-associated gut bacterium regulates the expression of pathogenic miRNAs during neuroinflammation. Specifically, we examined the relationship between *A. muciniphila* expansion and miR-21 induction during EAE and evaluated the contribution of this pathway to disease pathogenesis. Our findings identify a microbiota-dependent mechanism linking *A. muciniphila* to miR-21 production and provide evidence that microbiota-driven inflammatory miRNAs participate in gut-brain axis communication during autoimmune neuroinflammation.

## Method

### Mice

According to the National Institute of Neuroscience Animal Care Guidelines, C57BL/6, GF, and RAG KO mice were housed, fed, and maintained in a laboratory animal facility. All animal experiments were performed in accordance with institutional guidelines for the care and use of laboratory animals. All protocols were approved by the Committee on Ethics of Animal Experiments at the National Institute of Neuroscience, Tokyo, Japan.

### Antibiotics

To investigate the role of the gut microbiome in miRNA effects, mice were administered a mixture of antibiotics to deplete the gut microbiome. The antibiotic mixture, comprising kanamycin (1 mg/mL), vancomycin (0.5 mg/mL), and neomycin (1 mg/mL) (Wako Pure Chemical Industries, Ltd., Osaka, Japan) in 200 μL nuclease-free water, was administered via oral gavage, seven days before immunization, and was continued until the date of the experiment. Bacterial depletion was confirmed by measuring fecal bacterial DNA concentration.

### Induction and scoring of EAE

EAE was induced as previously reported [31]. Briefly, female mice at 6–8 weeks of age were injected subcutaneously with 100 mg of MOG35–55 peptides (synthesized at the Toray Research Center, Japan). The peptide mixture was emulsified using complete Freund’s adjuvant (Difco, Franklin Lakes, NJ, USA) containing 1 mg of heat-killed *Mycobacterium tuberculosis* H37RA. Intraperitoneal (i.p.) injections of pertussis toxin (100 ng; List Biological Laboratories, Campbell, CA, USA) were delivered on the day of immunization and repeated 48 hours later. Progression of neurological deficits was quantified daily based on a 5-point scale: 0 = asymptomatic; 0.5 = tail weakness; 1 = partial tail paralysis; 1.5 = severe tail paralysis; 2 = total tail flaccidity; 2.5 = flaccid tail coupled with hind limb weakness; 3 = partial hind limb paralysis; 3.5 = severe hind limb paralysis; 4 = total paralysis of the hind limbs; 4.5 = paralysis extending to both hind and forelimbs; 5 = death.

### Cell isolation

Single-cell suspensions of splenocytes were prepared by mechanical disruption of the spleen as previously reported [31]. CNS leukocytes were obtained by flushing spinal cords with PBS and harvesting brains from the skull. Minced brain and spinal cord samples underwent enzymatic digestion for 40 min at 37 °C using 1.4 mg/mL Collagenase H and 100 µg/mL DNase I (Roche) in RPMI. Suspensions were passed through a 70 μm strainer, and cells were enriched via a 37%/70% discontinuous Percoll gradient (GE Healthcare Life Sciences, Tokyo, Japan). Lamina propria lymphocytes from the small intestine (SI) and large intestine (LI) were prepared as previously described [32]. In brief, epithelial cells were removed from colonic tissues by incubating in HBSS (Wako) containing 1 mM dithiothreitol and 20 mM EDTA at 37 °C for 20 min. The tissue remnants were minced and digested for 30 min at 37 °C in RPMI 1640 (Sigma-Aldrich) supplemented with 12.5 mM HEPES (pH 7.2), 2% FCS, antibiotics (100 U/mL penicillin, 100 μg/mL streptomycin), 0.5 mg/mL Collagenase D (Wako), and 0.5 mg/mL DNase I (Roche). Following filtration, cells were washed in RPMI containing 2% FCS and separated on a Percoll gradient. Spleen and mesenteric lymph node single-cell suspensions were prepared via routine mechanical disruption.

### Flow cytometry

Cells were stained for fluorescence analysis as previously reported [31]. Briefly, to prevent nonspecific Fc receptor interactions, we pretreated isolated cells with an anti-mouse CD16/CD32 blocking antibody (BioLegend) before antibody incubation. We then performed surface antigen staining on ice for 30 min in PBS containing 5% FCS. Dead cells were eliminated from the gating strategy by labeling samples with the Aqua Live/Dead fixable dye (Invitrogen). Monoclonal antibodies against CD4, CD8, CD11b, CD11c, CD19, CD45, and TCRβ were purchased from BD, BioLegend, or eBioscience. For internal cytokine tracking, leukocytes underwent a 4-hour stimulation with phorbol 12-myristate 13-acetate and ionomycin prior to flow cytometric analysis. We acquired all sample data on a BD FACSCanto II system using FACS Diva software, and evaluated the final population metrics with FlowJo V10.7 (Tree Star).

### Cell sorting

T cells were isolated from splenocytes using a CD4^+^ T cell MACS isolation kit with an AutoMACS separator, according to the manufacturer’s instructions (Miltenyi Biotec, Bergisch Gladbach, Germany), as previously described [31]. For naive CD4^+^ T cells, L/D, CD45, TCRβ, CD25, CD62L, and CD44 were used, and cells were sorted on a FACS Aria IIu (BD). To isolate endothelial cells, we used a gentleMACS Octo Dissociator (Miltenyi Biotec) according to the instructions in the Adult Brain Dissociation Kit manual (Miltenyi Biotec). The obtained single-cell suspension was then subjected to endothelial purification using the MACS endothelial isolation kit (Miltenyi Biotec), selecting for CD45- and CD31+ cells.

### T-cell differentiation and cytokine analysis

For in vitro T-cell differentiation, naive CD4^+^CD62L^hi^CD44^lo^ CD25- T cells from C57BL/6 mice were sorted using flow cytometry and activated with anti-CD3 (2 μg/mL) and anti-CD28 (2 μg/mL) antibodies. For Th17 conditioned medium, cells were differentiated in the presence of TGF-beta (3 ng/mL), IL6 (20 ng/mL), anti-IFNG (10 µg/ml), and anti-IL4 (10 µg/mL). For Th1-conditioned medium, cells were differentiated in the presence of IL-12 (10 ng/mL) and anti-IL-4 (10 µg/mL) antibodies. After 96 h, the supernatant was collected, and cytokines were analyzed using ELISA.

### CD4^+^ T cell transfer

CD4^+^ T cells were prepared from the spleen and inguinal lymph nodes on day 9 of MOG immunization of GFP^+^ C57BL/6 mice using a CD4^+^ T Cell Isolation Kit (Miltenyi Biotec) (purity >95%). The cells were cultured in 24-well plates at a density of 1 × 10^6 cells/well in the presence of 20 μg/mL MOG peptide. On day 4, cells in the supernatant were collected and stimulated overnight with 2 μg/mL anti-CD3 and 1 μg/mL anti-CD28 antibodies. The cells were collected and washed with 1× PBS. T cells were transferred to C57BL/6 mice along with pertussis toxin on days 0 and 2.

### Immunoblotting

Primary endothelial cells cultured in the presence of miR-21 mimics and anti-miR-21 oligonucleotides were trypsinized and homogenized in RIPA buffer (Thermo Fisher Scientific, MA, USA) supplemented with a 1× protease inhibitor cocktail (Roche Applied Science). Equal amounts of protein (25 μg) were resolved using 12% polyacrylamide gel electrophoresis. Proteins were transferred onto a PVDF membrane, and immunoblotting was performed using TIMP3 Polyclonal Antibody (Cat no PA5-116049, Thermo Fisher Scientific, USA), ADAM-17 monoclonal antibody (Cat no. 703077, Thermo Fisher Scientific), and GAPDH (Cat no sc-32233, Santa Cruz, USA). HRP-conjugated polyclonal secondary antibodies against mouse IgG (cat no P0260, Dako, Denmark) were used. Images were captured using GE Image Quanta LAS500.

### Microarray analysis of miRNA

As previously done [11]. We extracted total RNA from mouse serum utilizing the commercial miRNeasy Serum/Plasma Advanced Kit (Qiagen, Germany) following the standard protocol. To perform a comprehensive overview of miRNA abundance, the extracted RNA was hybridized to the 3D-Gene Human miRNA Oligo Chip ver. 20 (Toray, Tokyo, Japan), enabling the simultaneous interrogation of roughly 2500 miRNA targets. Sample labeling was completed using a 3D-Gene miRNA labeling kit (Toray) prior to chip incubation. Finally, we scanned the microarray slides and quantified the resulting hybridization signals using a 3D-Gene Scanner (Toray).

### Isolation, quantification, and labeling of exosomes

Exosomes are isolated using a previously described technique [11, 33]. We isolated mouse plasma by centrifuging whole blood at 10,000 × *g* for 30 min at 4 °C to deplete intact cells, debris, and macro-EVs. We then extracted exosomes using the MagCapture™ Exosome Isolation Kit PS (Wako Pure Chemical Industries, Japan) according to the manual. To isolate exosomes from cell culture configurations, we applied a differential centrifugation pre-treatment to 15–20 mL of medium (300 × *g* for 5 min, 1,200 × *g* for 20 min, and 10,000 × *g* for 30 min at 4 °C) to clear cellular components and debris. This processed medium was concentrated down to 1 mL via a Vivaspin 20-100K device (Cat#: VS2041, Sartorius, UK) before applying the MagCapture™ matrix. We determined exosome concentrations using the FluoroCet Exosome Quantitation Kit (System Biosciences, CA, USA). For tracking assays, we labeled the vesicle membranes with the ExoGlow™-Membrane EV Labeling Kit (System Biosciences) and measured the resulting signal output on a GloMax® luminometer/fluorometer (Promega, WI, USA).

### Exosome tracking

Exosomes isolated from EAE and WT mice were labelled as described above using a standard kit. The concentration of labelled exosomes was normalized based on their fluorescence expression. Subsequently, an equal number of exosomes was suspended in 1× PBS and injected into the mice through the tail vein. After 1 h of transfer, the mice were sacrificed, and tissues were collected and homogenized in RIPA buffer using a tissue disrupter. The homogenized samples were centrifuged, and the supernatant was collected. Fluorescence was measured using GloMax^®^ (Promega).

### Immunohistochemistry

In brief, 1 h after injecting fluorescently labelled exosomes, the brain and spinal cord were dissected, and cryosections were prepared using a Cryostar NX70 cryostat (ThermoFisher Scientific, UAS) and fixed in 4% paraformaldehyde at 25°C. The sections were then processed using microwave irradiation and defatted with 0.2% Triton X-100. The sections were blocked in 3% BSA at 25°C. Sections were stained with Cyanine5 conjugated PECAM-1(Biolegend, USA) and DAPI (Molecular probes, USA), later mounted with Fluor mount (Southern Biotech, Birmingham, AL, USA) and imaged with a BZ-X710 automated microscope using the BZ-X Viewer software (Keyence).

### Isolation and quantification of exosomal and cellular miRNAs

Total RNA was extracted from the plasma, and purified exosomes were obtained from the same amount of plasma using the Plasma/Serum Circulating and Exosomal RNA Purification Kit (51000, Norgen Biotek Corp., ON, Canada). Five picograms of celmiR-39 mimic (Applied Biosystems, CA, USA) were added as an external control. RNA was subjected to complementary DNA (cDNA) synthesis using a TaqMan MicroRNA Reverse Transcription Kit (4366597, Applied Biosystems, CA, USA), and PCR was performed using the TaqMan Fast Advanced Master Mix (ThermoFisher Scientific) and StepOnePlus real-time PCR system (ThermoFisher Scientific). Data for each miRNA were analyzed using the delta-delta Ct method with celmiR-39 data as external control and normalized to the amount of applied RNA to 1 The amount of total RNA was determined using the QuantiFluor RNA System (Promega). To detect cellular RNA, an miRNeasy kit (Qiagen, Venlo, Netherlands) was used in the extraction step. The miRNAs were then analyzed using the method described above. For the analysis of mRNA, extracted RNA was subjected to cDNA synthesis using PrimeScript RT Master Mix (TAKARA, Shiga, Japan), and PCR was performed using the TaqMan Fast Advanced Master Mix and StepOnePlus real-time PCR system. Data were normalized to the expression of the exogenous RNA control.

### Cell culture

STC-1 (ATCC, USA), HEK-Blue™ TLR4, and HEK-Blue™ TLR2 (InvivoGen, USA) cells were maintained in Dulbecco’s modified Eagle medium (Gibco, NY, USA), supplemented with 10% fetal bovine serum (FBS, Gibco) and penicillin-streptomycin antibiotics (Gibco). Primary endothelial cells isolated from C57BL/6 mice using a kit for the isolation and cultivation of endothelial cells from adult brains (MACS; Miltenyi Biotec) were maintained in EBM-2 basal medium (Lonza, MD, USA) supplemented with 10% FBS (Gibco) and 1 U/mL antibiotic-antimycotic (Gibco).

### TLR reporter assay

TLR reporter cell lines (InvivoGen) were cultured in DMEM containing 10% FBS and 1× antibiotics. At 80% confluency, HEK-Blue™ TLR4 and TLR2 reporter cells were harvested using a scraper and resuspended in HEK-Blue™ Detection medium (InvivoGen). The resuspended cells (5 × 10^4^ cells/well; 180 μL) were then incubated with 20 μL heat-inactivated (by incubation at 60 °C for 30 min) *A. muciniphila* and *Lactobacillus reuteri* or with purchased purified *Akkermansia muciniphila* LPS (Sigma Aldrich) and CpG (InvivoGen) in a 96-well plate. TLR2 and TLR4 inhibitors were added 3 h before the stimulants were added for the inhibitor assay. After 20 h of culture, the secreted embryonic alkaline phosphate (SEAP) level was measured at 620 nm.

### TLR stimulation assay

Approximately 5 × 10^5^ STC-1 cells were cultured in a 25 cm^2^ flask in DMEM with 10% EXO-free FBS (ThermoFisher Scientific). After reaching confluence, cells were treated with purchased purified *A. muciniphila* LPS and CpG at 1, 10, and 100 ng/mL concentrations for 24 h. The culture medium was collected for quantification of exosome and cell-free miRNAs.

### Intestinal permeability assay

On day 7 postimmunization, the mice were fasted for 6 hours before testing. Thereafter, approximately 50 μL of blood was collected by tail nicking, and then 150 μL/mouse of 80 mg/mL FITC-dextran (4 kDa; Sigma) in sterile 1× PBS was administered by oral gavage. Mice were euthanized, and blood was sampled 4 h after the gavage. The pre- and post-test plasma samples were diluted 1:10 in 1× PBS and transferred to a black, opaque-bottom 96-well plate. The concentration of FITC-dextran was determined by measuring the fluorescence at 530 nm after excitation at 485 nm.

### Systemic administration of miRNAs and antisense-oligonucleotides

For systemic administration of synthetic miRNAs and antisense-oligonucleotides previously establish technique was used [34]. Briefly, 15 μM miRNA–atelocollagen mixture was prepared using an AteloGene Systemic Use kit (KOKEN), according to the manufacturer’s instructions, and 200 μL of the mixture was intravenously administered per mouse via the tail vein. DC-CHOL/DOPE cationic liposomes (FormuMax Scientific Inc., Sunnyvale, CA, USA) were used as an alternative. A 15 μM miRNA-DC-CHOL/DOPE complex was prepared in 200 μL PBS and intravenously administered from the tail vein to each mouse. For anti-miR oligo treatment, anti-miR-21, anti-miR-223, anti-miR-146, and anti-Cel-miR-39 controls (30 μg/mouse) were administered i.v. to MOG-immunized mice on days 0, 7, and 14 after immunization. For the passive transfer experiment, Mimic miR-21, anti-miR-21, and control miR were administered on days 0, 2, and 4 after cell transfer.

### Transfection and reporter assay

As previously described [34], the day before transfection, cells were trypsinized and seeded in 96-well culture plates (0.5 × 10^4^ cells/well). The constructed Goclone® reporter vectors are transfection-ready and were transfected into cells with synthetic miRNA mimics (20 nM final concentration) using Lipofectamine 2000 (Thermo Fisher Scientific) according to the manufacturer’s instructions. After 24 h of transfection, the cells were lysed, and luciferase activity from the luminescent reporter gene (RenSP) was measured using the Lightswitch^TM^ Assay Kit (Switchgear Genomics) according to the manufacturer’s protocol. Fold changes were calculated and compared with those in control cells.

### RNA and DNA oligonucleotides

The RNA and DNA oligonucleotides used in this study were synthesized by Hokkaido System Science (Sapporo, Japan), order number 41033160. The sequences of the synthesized oligonucleotides are described below. The mirVANA^TM^ microRNA mimics were purchased from Thermo Scientifics (USA).

### RNA mimics

mus-miR-21a-5p_S - UAG CUU AUC AGA CUG AUG UUG A

mus-miR-21a-5p_AS - UCA ACA UCA GUC UGA UGG GCU GUC

si Cont_S - UUC UCC GAA CGU GUC ACG UUU

si Cont_AS - ACG UGA CAC GUU CGG AGA AUU

### DNA oligonucleotide (Anti-sense-miRNAs)

AntiS miR-21 – TCA ACA TCA GTC TGA TAA GCT A

AntiS miR-146–AAC CCA TGG AAT TCA GTT CTC A

AntiS miR-223–TGG GGT ATT TGA CAA ACT GAC A

AntiS CelmiR-39 – CAA GCT GAT TTA CAC CCG GTG A

### Isolation and culture of bacteria

To isolate *L. reuteri*, feces from wild-type C57BL/6 mice were homogenized and diluted with PBS. The dilutions were plated on YCFA agar plates and incubated in an anaerobic chamber for two days as previously shown [32]. The 16S rRNA genes from individual colonies were amplified using the 27F and 1492R primers. We then performed Sanger sequencing with the 27F and 519R primers to identify colonies corresponding to *L*. *reuteri. Akkermansia muciniphila* JCM 33894 was procured from the RIKEN BRC Microbe Division. These bacteria were grown in GAM broth and stored in 20% glycerol at −80 °C.

### Bacterial colonization into mice

The type strain of *Akkermansia muciniphila- JCM 30893* was grown in mucin added GAM broth. *Lactobacillus reuteri* strains were isolated from day 7 EAE mice grown on MRS agar further expanded in mucin added GAM broth. *L. reuteri* was administered as a control bacterium. Live bacteria (OD600 0.40, 200 μL) or vehicle control of mucin broth were delivered to 7-week-old female C57BL6J mice (n = 5/group) by oral gavage on day -9 before disease induction. Antibiotic administration was performed for 7 days, and 2, 2-day wash gap was given. Further induction of Experimental autoimmune encephalomyelitis and neurological deficits were evaluated on a scale from 0 to 5, as mentioned above. For mono colonization of GF mice, *Akkermansia muciniphila- JCM 30893* and *Lactobacillus reuteri* expanded in Mucin added GAM broth were administrated twice a week for 3 weeks and experiments were performed after 6 days of last administration. GF mice were induced EAE as mention above where as purified *A muciniphila* LPS is administrated on day 3, 6 and 9 post immunization.

### Shotgun metagenomic sequencing

Sequencing libraries were constructed from metagenomic DNA using an Illumina DNA PCR-Free Prep, Tagmentation kit (Illumina, San Diego, CA, USA) and an IDT for Illumina DNA/RNA Unique Dual Indexes (Illumina), according to the manufacturer’s instructions. Each library was size-selected and cleaned using Illumina Purification Beads. The pooled sample comprised 35 metagenomic libraries and was sequenced on an Illumina NovaSeq 6000 using an SP Reagent Kit v1.5 with 150 bp paired-end sequencing, yielding an average of 23 million read pairs per library. Shotgun metagenomic sequencing data are available from DDBJ DRA under the BioProject ID PRJDB 17480.

### Shotgun metagenomic sequence analysis

Short read pairs were quality-filtered using fastp version 0.2026 with the parameter “-n 1 -l 70.” [35]. VITCOMIC2 was used to assign taxa to the sequence reads [36]. Quality-filtered reads were assembled using the MEGAHIT version 1.2.927 [37]. The assembled contigs were subjected to metagenomic binning using the MetaBAT2 version 2.15 [38]. All bins were quality checked (completeness >50% and contamination rate <10%) [39] and taxon assigned using the CheckM version 1.13 and GTDB-Tk version 1.0.2 [40], respectively. Protein-coding genes were predicted using the Prodigal version 2.6.3 in the metagenomic mode. [41]. To quantify gene abundance, reads were mapped to contigs using the bwa-mem version 0.7.17 [42]. The transcripts per kilobase million (TPM) for each gene were calculated from the read-mapping results. [43]. Gene functions were annotated using the KEGG orthology. [44] from a protein vs. protein sequence similarity search using the DIAMOND version 2.0.9, with the parameters “--sensitive --value 1e-16 --id 40 - -strand both.” [45]. The *A. muciniphila* metagenome-assembled genomes and the complete genome of *A. muciniphila* JCM 30893 were orthologous clustered using the OrthoFinder2 version 2.5.2 [46]

### Quantification and statistical analysis

Statistical analyses were performed using Prism 10.0 (GraphPad) software. Unless mentioned, the Unpaired two-tailed Student’s t-test and the Unpaired two-tailed Mann-Whitney test were used to calculate the *P* values. The exact *P* values were mentioned in the figures, and P values <0.05 were statistically significant. Data are presented as mean ± SD. Principal Component (PCA) analysis was performed using the Orange data mining tool.

### Data and code availability

All data required to evaluate the conclusions in the paper are presented in the paper and/or Supplementary Materials. Shotgun metagenomic sequencing data are available from DDBJ DRA under the BioProject ID PRJDB 17480. The Gene Expression Omnibus (GEO) accession number for the miRNA expression data used in this study is GSE255469.

## Results

### Gut microbiota regulate circulating miRNA expression during EAE

To investigate whether gut microbiota regulate circulating miRNAs during autoimmune neuroinflammation, we first established a microbiota dysbiosis model by administering a broad-spectrum antibiotic cocktail beginning 7 days before MOG35–55 immunization and continuing throughout the experiment (Figure S1A). Antibiotic-treated mice (ABX-EAE) developed significantly less severe disease than untreated EAE mice (Figure 1A). In contrast, four days of antibiotic treatment before immunization failed to alter disease progression, indicating that sustained depletion of the microbiota was required to confer protection (Figure S1B). Consistent with successful microbiota disruption, fecal microbial DNA content was markedly reduced on day 7 after immunization and partially recovered by day 14 (Figure S1D). Microbiome analysis further demonstrated altered microbial diversity and community composition in both EAE and ABX-EAE mice compared with normal controls (Figure 1B and Figure S1E). Intestinal permeability was increased on days 7 and 14 after immunization in both EAE and ABX-EAE mice (Figure S1C). We next examined whether microbiota disruption affected circulating miRNA profiles during EAE. Microarray analysis of serum collected at day 14 post-immunization revealed distinct miRNA expression patterns among control, EAE, and ABX-EAE mice (Figure 1C). Principal component analysis demonstrated clear separation of EAE samples from both control and ABX-EAE groups, indicating disease-associated changes in circulating miRNA composition (Figure 1D). Among the differentially expressed miRNAs, miR-21a-5p, miR-122-5p, miR-146a-5p, and miR-223-3p were markedly elevated in EAE mice but remained suppressed in ABX-EAE mice and controls (Figure 1D), suggesting that induction of these circulating miRNAs depends on the presence of an intact microbiota.

**Figure. 1:**
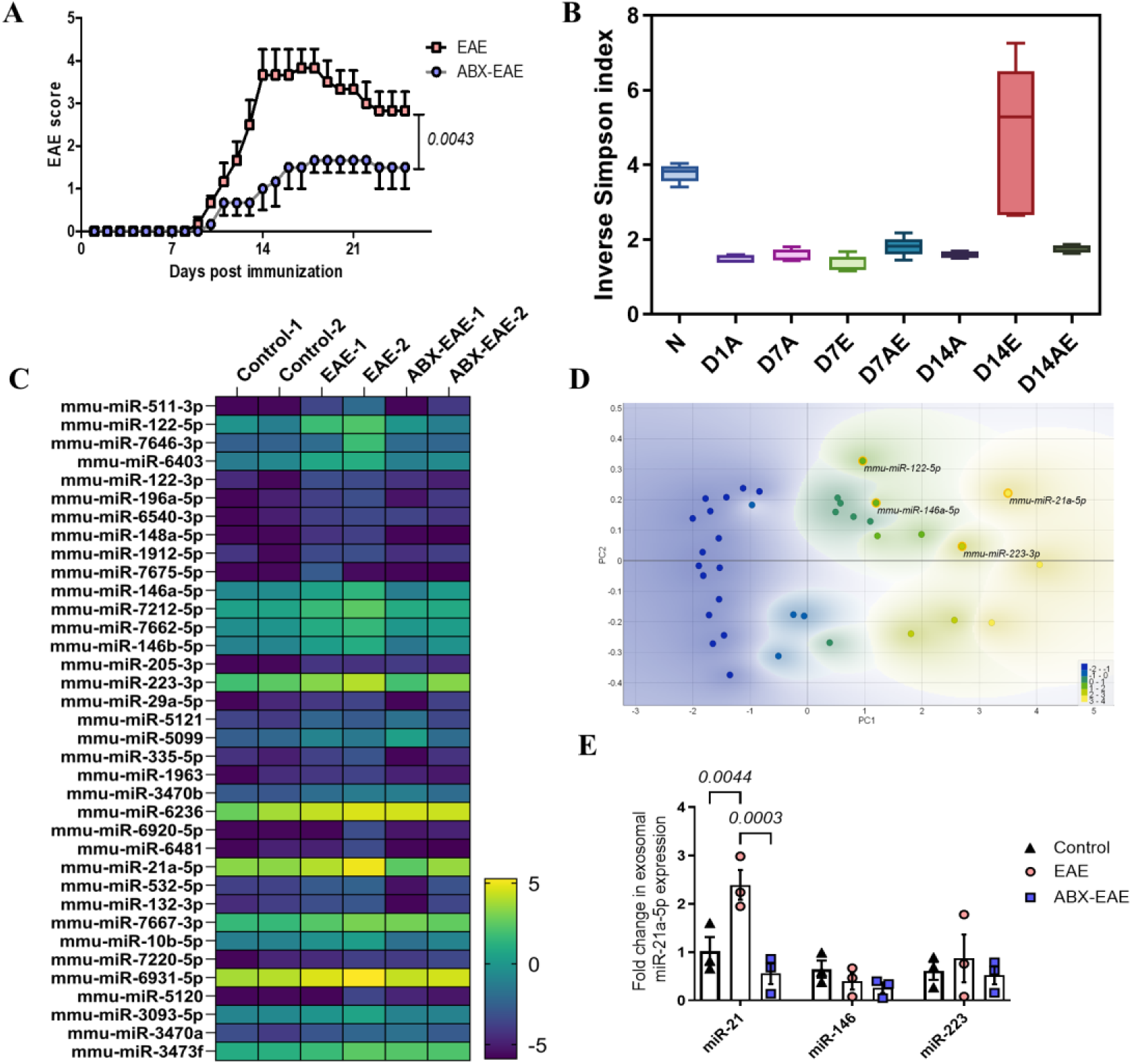
Significant change in gut microbiota and blood-circulating miRNAs between antibiotic-treated and untreated EAE mice. **A.** The mean clinical score of the MOG_35-55_-induced EAE between antibiotic-treated (ABX-EAE) and non-antibiotic-treated EAE mice. Mice were orally administered with an antibiotic cocktail (ABX) every day starting from one week before immunization and continued till the date of the experiment (n=5 individual data). **B.** Inverse Simpson index of fecal microbiome. To analyse the difference in the microbial diversity,16S rRNA analysis was performed in the feces collected from normal (N), EAE (E), ABX-EAE (AE), and ABX-treated non-EAE (A) mice at Day 7 (D7) and Day 14 (D14) after immunization (n=5 individual data). **C**. Microarray analysis of blood circulating miRNAs. Micro RNAs were isolated from the freshly collected serum of control, EAE, and ABX-EAE mice on day 14 after immunization and subjected to microarray analysis. **D.** Principal component analysis on microarray data. Circulating miRNAs significantly differentially expressed between EAE and control mice were represented using PCA analysis. PC1 represents control, and PC2 represents EAE. Overexpressed miRNAs are categorized with the yellow outline. **E.** qRT-PCR analysis of exosomal miRNAs. Blood circulating exosomes were isolated from freshly collected serum of control, EAE and ABX-EAE mice on day 14 after immunization (n=3 individual data). Exosomal miRNAs were isolated by MagCapture(TM) kit, and exosomal miR-21a-5p, miR-146a-5p, and miR-223-3p were examined by qRT-PCR. Figure C data represented in duplicates. Error bars represent the mean ±SEM values. In Figure A, a one-sample T test was used, and in Figure E, a 2-way ANOVA was used to calculate the *P* values.

To determine whether induction of these miRNAs was associated with antigen-specific autoimmunity, we analyzed their expression following immunization with complete Freund’s adjuvant (CFA) alone. Among the four candidate miRNAs, only miR-122 was induced in the absence of MOG35–55, whereas miR-21, miR-146a, and miR-223 remained unchanged (Figure S1F). Longitudinal analysis further demonstrated that miR-21, miR-146a, and miR-223 were elevated during both the early (day 7) and progressive (day 14) phases of EAE (Figure S1G), indicating sustained induction throughout disease development.

Because circulating miRNAs can be transported within extracellular vesicles, we next quantified serum exosomes and exosome-associated miRNAs. Exosome numbers were increased in both EAE and ABX-EAE mice compared with normal controls (Figure S1H). Analysis of exosomal miRNAs revealed significantly elevated miR-21 expression in EAE mice, whereas this increase was attenuated in ABX-EAE mice (Figure 1E). Changes in exosomal miR-146a and miR-223 were less pronounced. Collectively, these findings demonstrate that disruption of the gut microbiota attenuates EAE severity and suppresses the induction of disease-associated circulating and exosomal miRNAs, identifying miR-21 as a prominent microbiota-dependent candidate associated with EAE.

### Microbiota-dependent miR-21 promotes EAE pathogenesis

To determine whether microbiota-regulated circulating miRNAs contribute to EAE pathogenesis, we inhibited miR-21, miR-146a, and miR-223 in vivo using antisense oligonucleotides administered on days 0, 7, and 14 after MOG35–55 immunization (Figure 2A). Inhibition of all three miRNAs attenuated EAE severity, with anti-miR-21 producing the strongest reduction in clinical scores (Figure 2A). Consistent with this finding, anti-miR-21 significantly reduced T-cell infiltration into the CNS (Figure 2B), identifying miR-21 as the most prominent disease-associated candidate. Because miR-21 expression was suppressed in ABX-EAE mice, we next examined its dependence on the gut microbiota. Circulating miR-21 levels were significantly higher in specific-pathogen-free (SPF) mice than in germ-free (GF) mice and were robustly induced following EAE induction only under SPF conditions (Figure 2C). In contrast, EAE induction failed to increase circulating miR-21 in GF mice (Figure 2C), indicating that microbiota-derived signals are required for its induction.

**Figure. 2:**
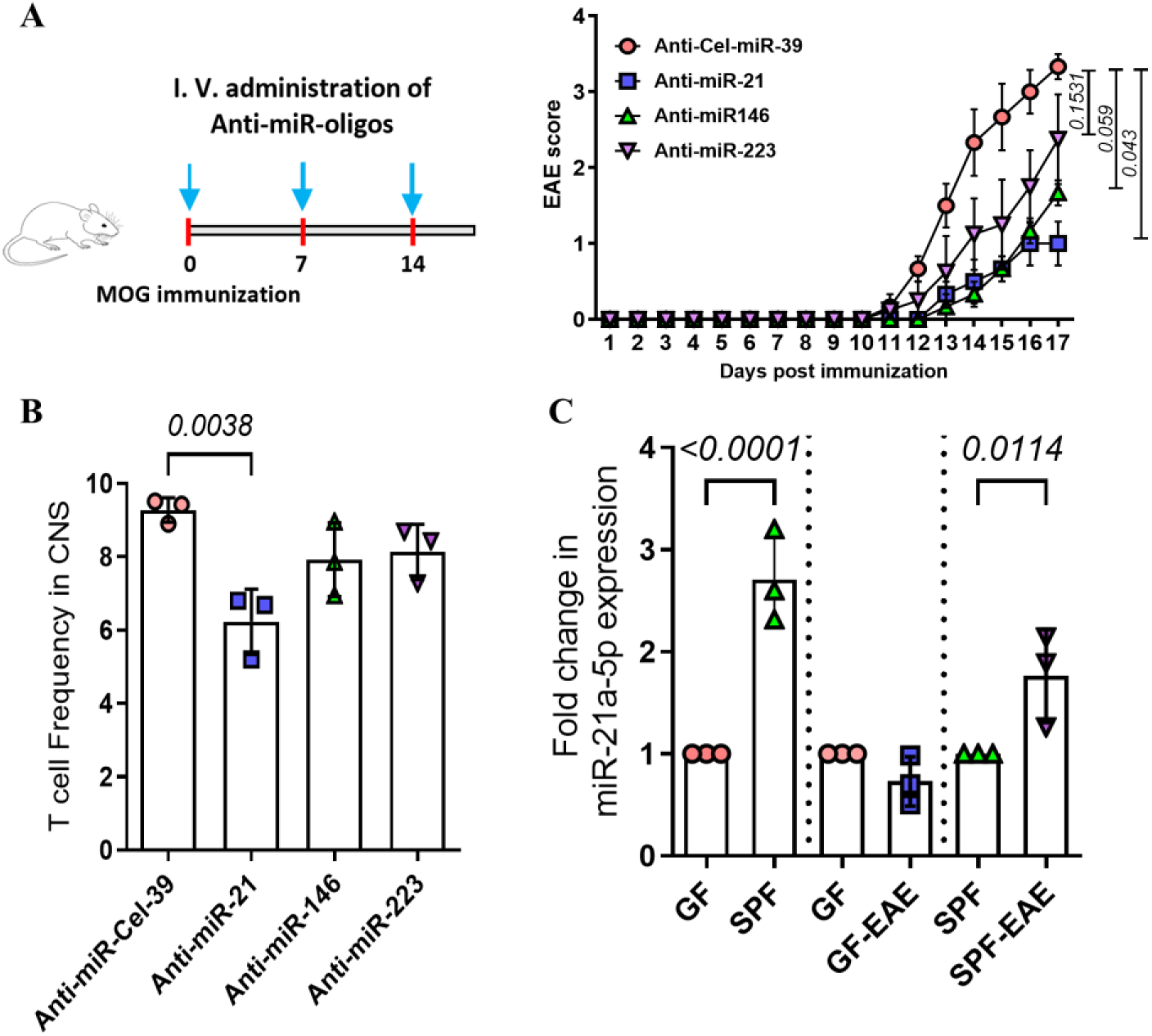
Blood-circulating miR-21 related to EAE via gut microbiota. **A.** Experimental schematic and the mean clinical EAE score of EAE mice treated with antisense oligonucleotides of miRNAs. To suppress the overexpressed blood circulating miRNAs in EAE, EAE mice were treated with intravenous injection of antisense oligonucleotides-atelocollagen mixtures of miR-21, miR-146, and miR-223 on days 0, 7, and 14 post-immunization. Clinical EAE score was recorded till the date of the experiment (n=3 independent mice). **B.** Frequency of T cells entering the CNS. Total T cell frequency in the CNS of mice treated with antisense oligonucleotides-atelocollagen mixtures of miR-21, miR-146, and miR-223 was analysed at day 18 post-immunization. **C.** qRT-PCR analysis of blood circulating miR-21 in germ-free and SPF mice. The level of circulating miR-21 in blood was quantified from freshly collected serum of germ-free and SPF mice, both without immunization and on day 14 after immunization (n=3 independent mice). Error bars represent the mean ± SEM. In Figure A, a two-tailed Mann-Whitney U test was used; in Figures B and C, ANOVA was used to calculate the *P* values.

To determine whether adaptive lymphocytes were the sole source of circulating miR-21, we analyzed Rag-knockout (Rag-KO) mice lacking mature T and B cells. Despite the absence of adaptive lymphocytes, EAE induction significantly increased circulating miR-21 levels in Rag-KO mice (Figure S2A), suggesting the contribution of additional cellular sources. Collectively, these findings identify miR-21 as a microbiota-dependent circulating miRNA that contributes to EAE severity.

### Expansion of *Akkermansia muciniphila* is associated with elevated circulating miR-21

Because circulating miR-21 expression depended on the gut microbiota, we next sought to identify bacterial taxa associated with its induction during EAE. Longitudinal 16S rRNA sequencing of fecal samples revealed marked shifts in microbial composition during disease progression (Figures 3A and S3A). At day 7 post-immunization, EAE mice exhibited increased abundance of several genera, including *Akkermansia*, *Olsenella*, *Bilophila*, and *Barnesiella*, together with reduced abundance of *Lactobacillus and Escherichia/Shigella* compared with ABX-EAE and control mice (Figures 3A, 3B, and S3A). Among these changes, *Akkermansia* showed the most prominent enrichment, accounting for approximately 70% of the bacterial population in day 7 EAE mice (Figure 3B).

**Figure. 3:**
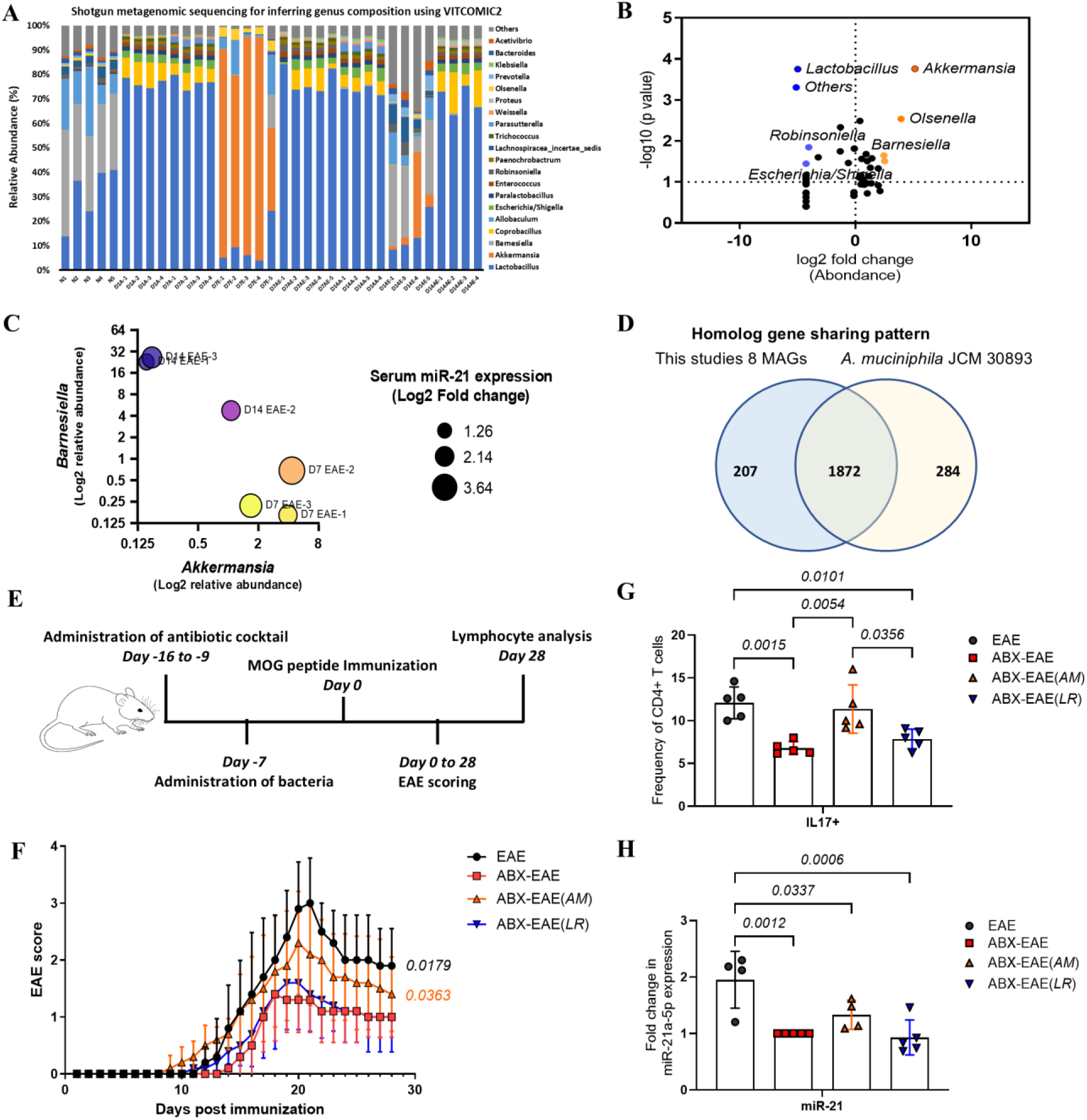
Increased *Akkermansia* and miR-21 in EAE. **A.** Fecal microbiome analysis. Shotgun metagenomic sequencing data of fecal samples collected from normal (N), dysbiosis (A), dysbiosis EAE (AE), and EAE (E) mice on days 7 and 14 using VITCOMIC 2 are exhibited. **B.** Differential relative abundance of bacterial taxa. The graph shows the abundance of bacteria on day 7 in EAE mice; the x-axis shows abundance, and the y-axis shows the P value. Red and blue indicate increases and decreases in relative abundance, respectively. **C.** Multiple variable analysis on miR-21 and bacteria abundance. The graph–shows the log2-fold change in miR-21 expression on day 7 relative to the abundance of *A. muciniphila* and *Barnesiella* (n=3 independent mice). **D.** Venn diagram of whole-genome sequence. The draft genome sequence of *A muciniphila* derived from EAE mice is shown, along with a known sequence of *A muciniphila* JCM 30893 (comparison based on OrthoFinder2). **E.** Experimental schematic. To confirm the role of bacteria in EAE induction, four types of treatments were performed. Antibiotics-treated mice were colonized with *Akkermansia muciniphila* JCM 30893 (ABX-EAE(*AM*)) and *Lactobacillus reuteri* (ABX-EAE(*LR*)) and immunized with MOG_35-55._ As a control, mice treated with or without antibiotics were subjected to EAE (ABX-EAE and EAE, respectively). **F.** EAE score. The mean EAE clinical score in ABX-EAE(*AM*), ABX-EAE(*LR*), ABX-EAE, and EAE was recorded (n=5 independent mice). **G.** Cytokine analysis: CD4+ T cells were isolated from post-immunized mice on day 28 and stimulated with PMA for 4h, and intracellular cytokine expression was examined (n=5 independent experiments). **H.** qRT-PCR analysis of blood circulating miR-21. Freshly collected serum from experimental mice was used to quantify circulating miR-21 on day 28 post-immunization. (n=5 independent experiments). Error bars represent mean ± SEM values. In F, a one-sample T test was used, and in G and H, an ordinary 2-way ANOVA was used to calculate the *P* values.

To determine whether specific bacterial taxa were associated with circulating miR-21 expression, we compared bacterial abundance with serum miR-21 levels. The relative abundance of *Akkermansia* positively correlated with circulating miR-21 expression during early EAE, whereas *Barnesiella* showed no comparable relationship (Figure 3C). To obtain species-level resolution, we subsequently performed shotgun metagenomic sequencing. This analysis identified the enriched *Akkermansia* species as *Akkermansia muciniphila* and the predominant *Lactobacillus* species in ABX-EAE mice as *Lactobacillus reuteri* (Figure 3A). Comparative genomic analysis further demonstrated substantial gene conservation between the EAE-derived *A. muciniphila* isolate and the reference strain *A. muciniphila* JCM 30893 (Figure 3D).

To assess the functional relevance of these bacterial species, ABX-treated mice were colonized with either *A. muciniphila* JCM 30893 or *L. reuteri* before EAE induction (Figure 3E). Colonization with *A. muciniphila* significantly increased EAE severity compared with ABX-EAE mice, whereas *L. reuteri* failed to restore disease activity (Figure 3F). Consistent with these clinical findings, *A. muciniphila*-colonized mice exhibited increased frequencies of IL-17⁺ CD4⁺ T cells in the CNS (Figure 3G). Serum miR-21 expression was also significantly elevated in *A. muciniphila*-colonized mice compared with ABX-EAE animals (Figure 3H). In contrast, monocolonization of germ-free mice with either *A. muciniphila* or *L. reuteri* in the absence of EAE did not significantly alter circulating miR-21 levels (Figure S3B). Collectively, these findings identify *A. muciniphila* as a bacterial species associated with increased miR-21 expression and enhanced neuroinflammatory responses during EAE.

### *Akkermansia muciniphila* LPS induces epithelial miR-21 through TLR2/4 signaling

To identify microbial functions associated with the expansion of *A. muciniphila* during EAE, we performed KEGG-based functional profiling of metagenomic datasets from EAE and ABX-EAE mice. Distinct distributions of KEGG orthologs and pathways were observed between groups (Figures S4A and S4B). In EAE mice, day 7 samples were enriched for several microbial pathways, including ribosomal function, bacterial secretion systems, and lipopolysaccharide (LPS) biosynthesis (Figure 4A). These functional changes coincided with the marked expansion of *A. muciniphila* during early EAE.

**Figure. 4:**
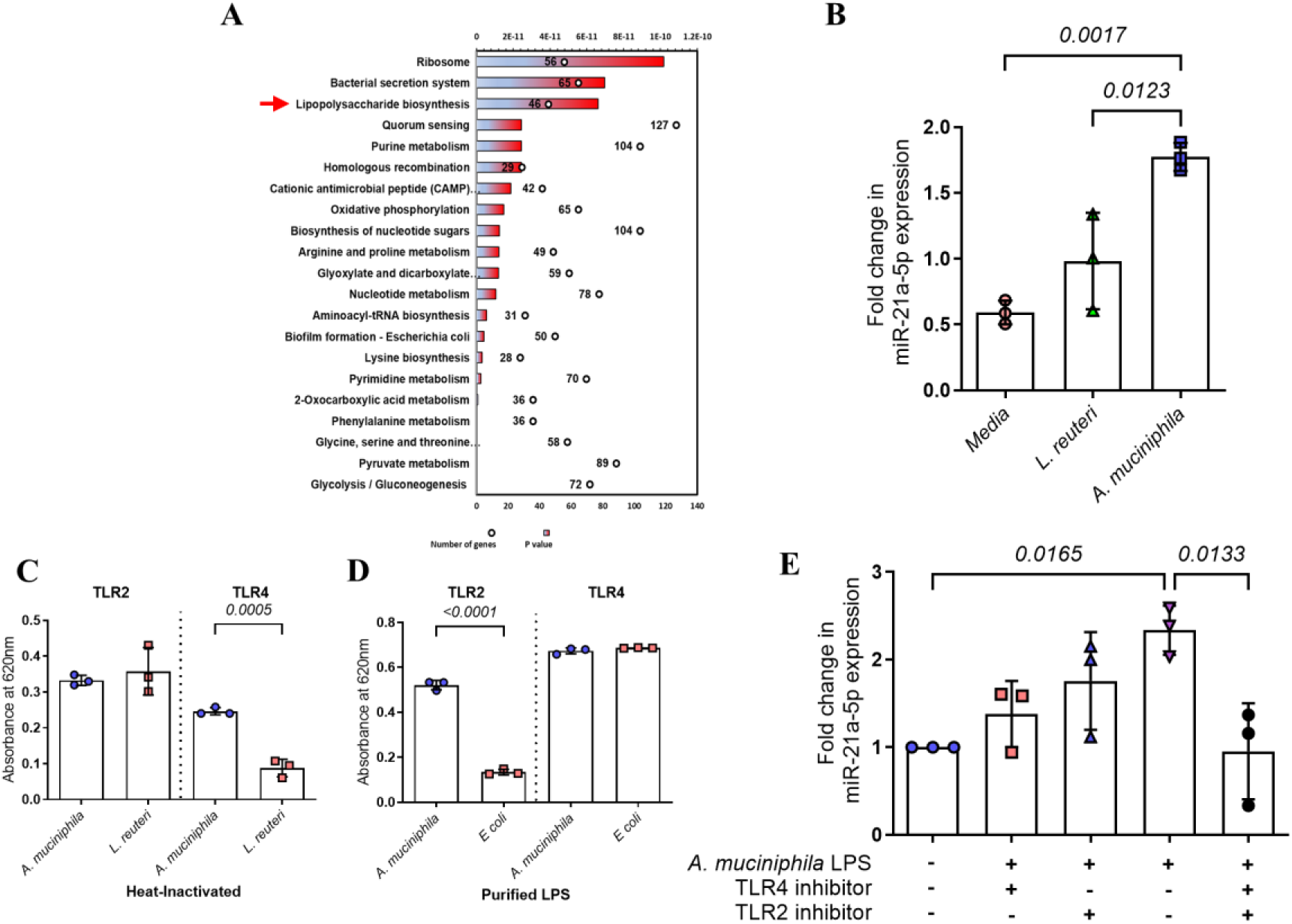
*A. muciniphila* LPS induces epithelial miR-21 through TLR. **A.** Microbial functional analysis in EAE. KEGG pathways enriched with KOs on day 7 EAE, listed based on the significant p-value. **B.** Analysis of miR-21 in gut epithelial cells. Intercellular (cell-free) miR-21 in the culture medium of STC-1 cells, a mouse-derived intestinal epithelial cell line, stimulated with heat-inactivated *A. muciniphila* and *L. reuteri*, was examined by qRT-PCR (n=3 independent determinations). **C.** TLR reporter cell assay. TLR2 and TLR4 reporter cell assays were performed with reporter cells individually stimulated by heat-inactivated *A. muciniphila* and *L. reuteri*. After 20 h of culture, the level of secreted embryonic alkaline phosphate (SEAP) was measured at 620 nm (n=3 independent determinations). **D.** TLR reporter cell assay. TLR2 and TLR4 reporter cell assays were performed with reporter cells stimulated by purified LPS of *A. muciniphila* and *E. coli*. After 20 h of culture, the level of secreted embryonic alkaline phosphate (SEAP) was measured at 620 nm (n=3 independent determinations). **E.** Analysis of miR-21 in gut epithelial cells. Cell-free miR-21 in the medium of STC-1 cells stimulated with purified *A. muciniphila* LPS in the presence or absence of TLR2 and TLR4 inhibitors was examined by qRT-PCR (n=3 independent determinations). Error bars represent the mean ±SEM values. In C and D, a two-tailed Mann-Whitney U test was used; in B and E, ANOVA was used to calculate *P* values.

Because epithelial miR-21 expression was elevated during EAE (Figure 5A) and intestinal barrier permeability increased during disease progression (Figure S1C), we next investigated whether *A. muciniphila* directly induces epithelial miR-21 expression. Stimulation of STC-1 intestinal epithelial cells with heat-inactivated *A. muciniphila* resulted in significantly greater miR-21 secretion than stimulation with *Lactobacillus reuteri* (Figure 4B). We therefore examined the contribution of *A. muciniphila* LPS. Purified *A. muciniphila* LPS induced dose-dependent increases in both cell-free and exosome-associated miR-21 secretion, whereas CpG stimulation produced substantially weaker responses (Figures S4C and S4D). Consistent with these findings, oral administration of purified *A. muciniphila* LPS to EAE-induced germ-free mice significantly increased circulating miR-21 levels compared with untreated GF-EAE mice (Figure S4E).

**Figure. 5:**
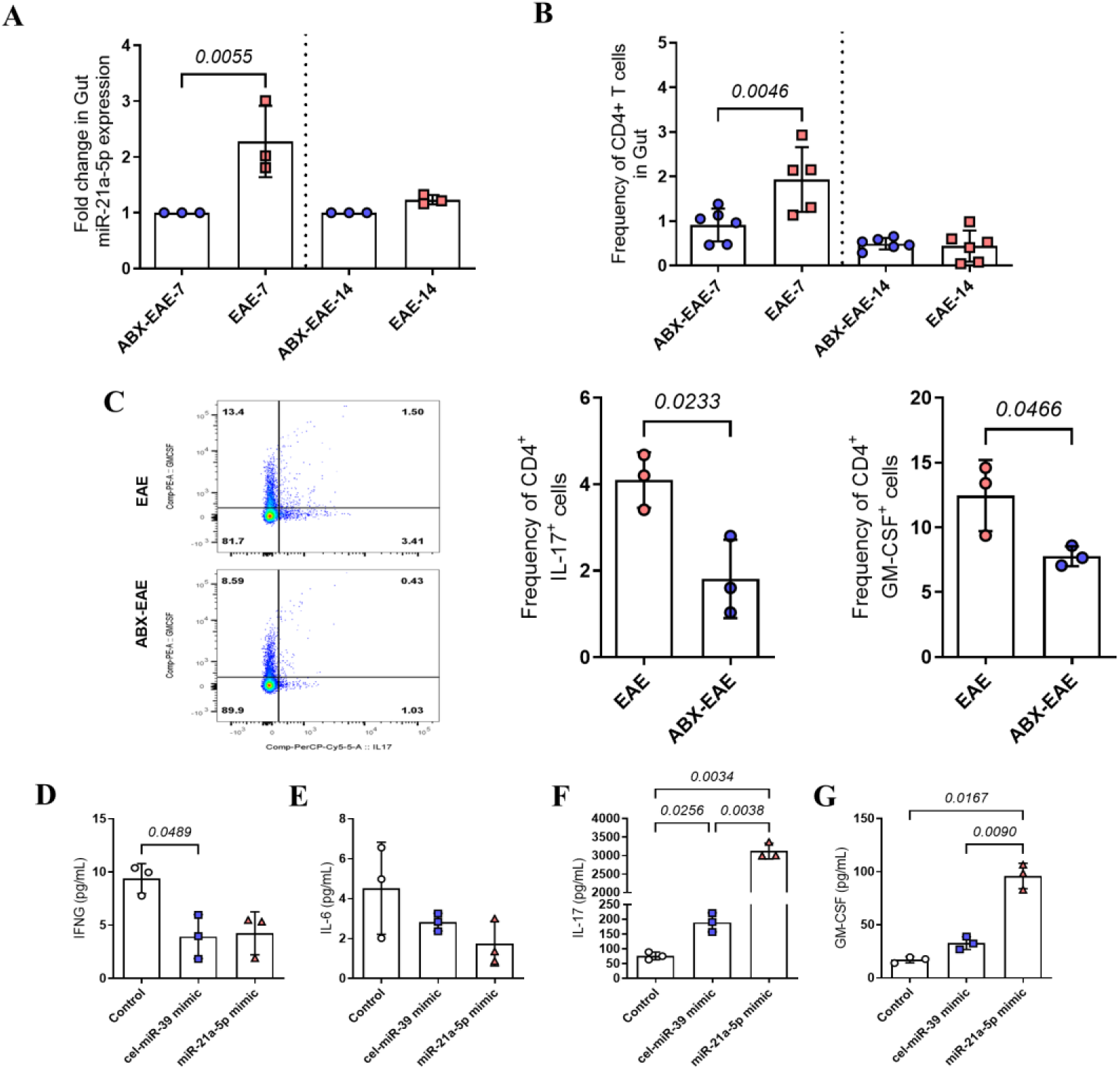
Epithelial miR-21 and T cells in the large intestine of EAE and ABX-EAE mice. **A.** qRT-PCR analysis of miR-21 expression in gut epithelial cells. Intracellular expression of miR-21 was examined in small and large intestine epithelial cells in EAE and ABX-EAE mice at day 7 post-immunization (n=3 independent experiments). **B.** CD4^+^ T cell screening in the gut. Frequency of CD4^+^ T cells was analyzed in the small (SI) and large (LI) intestines on days 7 and 14 in EAE and ABX-EAE (n=5 to 7 individual mice per experimental group). **C.** Intracellular cytokine expression in CD4^+^ cells derived from the gut. CD4+ cells from the colonic lamina propria of EAE and ABX-EAE mice on day 7 were stimulated with PMA for 4 hours, and the expression of intracellular cytokine levels was examined. (n=3 independent experiments). **D-G.** Effects of miR-21 on CD4^+^ T cell cytokine secretion. Synthetic miR-21 mimic (Mission® microRNA mimics) and cel-miR-39 (as a nonspecific control) were transfected into naive CD4^+^ cells in Th17 conditioned medium, and the cytokine secretion was measured after 96 h using ELISA (n = 3 independent determinations). Error bars represent the mean ± SEM values. In figures A-C a two-tailed Mann-Whitney U test was used, in figures D-G Brown-Forsythe and Welch ANOVA were used to calculate the *P* values.

To define the signaling pathways involved, we examined TLR activation using reporter cell assays. Heat-inactivated *A. muciniphila* preferentially activated TLR4, whereas *L. reuteri* exhibited minimal activity (Figure 4C). Purified *A. muciniphila* LPS activated both TLR2 and TLR4 reporter cells, confirming the atypical TLR-stimulatory properties of this LPS preparation (Figure 4D). Pharmacological inhibition of either TLR2 or TLR4 reduced LPS-induced miR-21 expression in STC-1 cells, while simultaneous activation of both pathways produced the highest miR-21 levels (Figure 4E). These findings indicate that *A. muciniphila* LPS promotes epithelial miR-21 production through combined TLR2/TLR4 signaling.

Because miR-21 was secreted in both cell-free and exosomal forms, we next examined the biological properties of epithelial-derived exosomes. Exosomes isolated from LPS-stimulated epithelial cells were fluorescently labeled and transferred into SPF recipient mice (Figure S4F). Compared with control or CpG-derived exosomes, LPS-induced exosomes showed enhanced accumulation within the brain and spinal cord (Figures S4G and S4H). Furthermore, exosomes derived from LPS-stimulated epithelial cells increased IL-17 and GM-CSF production from CD4⁺ T cells cultured under Th17-polarizing conditions (Figure S4I). Collectively, these findings identify *A. muciniphila* LPS as a potent inducer of epithelial miR-21 via TLR2/TLR4 signaling and demonstrate that epithelial-derived exosomes can disseminate inflammatory signals that enhance pathogenic T-cell responses.

### miR-21 enhances pathogenic Th17 responses during EAE

Commensal bacteria have been reported to induce miR-21 expression in intestinal epithelial cells [22]. We therefore examined epithelial miR-21 expression in EAE and ABX-EAE mice on days 7 and 14 post-immunization. Epithelial miR-21 expression was significantly elevated in day 7 EAE mice compared with ABX-EAE mice (Figure 5A), coinciding with increased accumulation of intestinal CD4⁺ T cells during the early phase of disease (Figure 5B). Consistent with this observation, CD4⁺ T cells isolated from the colonic lamina propria of day 7 EAE mice produced higher levels of IL-17 and GM-CSF following PMA/ionomycin stimulation than those from ABX-EAE mice (Figure 5C).

To investigate the potential role of miR-21 in regulating T-cell inflammatory responses, we performed gene ontology analysis, which identified positive associations between miR-21 and cytokine production pathways (Figure S5A). Bioinformatic analysis further identified a conserved miR-21 binding site within the 3′ untranslated region (UTR) of SMAD7 mRNA, and luciferase reporter assays confirmed direct interaction between miR-21 and the SMAD7 3′UTR (Figures S5B-C). We next examined the functional effects of miR-21 on CD4⁺ T cells. Naïve CD4⁺ T cells were transfected with synthetic miR-21 mimics and cultured under Th17-polarizing conditions. miR-21 overexpression significantly increased IL-17 and GM-CSF secretion, whereas IFN-γ and IL-6 production remained unchanged (Figures 5D-G). In contrast, miR-21 overexpression did not significantly alter cytokine production under Th1-polarizing conditions (Figure S5D). Collectively, these findings identify miR-21 as a positive regulator of Th17-associated inflammatory responses during EAE, potentially by suppressing SMAD7.

### Circulating miR-21 promotes CNS infiltration through a TIMP3–ADAM17 pathway

Because circulating exosomes are stable and can access the central nervous system (CNS), we first examined their tissue distribution during EAE. Exosomes isolated from specific pathogen-free (SPF) and EAE mice at peak disease were fluorescently labeled and intravenously transferred into SPF recipients (Figure S6A). Labeled exosomes distributed broadly across peripheral tissues and were readily detected in the brain and spinal cord (Figures S6B and S6C), indicating that circulating exosomes can access CNS tissues during neuroinflammation.

To determine whether circulating miR-21 influences CNS immune cell infiltration, MOG-reactive GFP⁺ CD4⁺ T cells were adoptively transferred into recipient mice, followed by administration of miR-21 mimics or anti-miR-21 oligonucleotides (Figure 6A). Mice receiving miR-21 mimics exhibited significantly increased accumulation of GFP⁺ CD4⁺ T cells within the CNS compared with mice receiving anti-miR-21 treatment (Figures 6B and 6C). Similarly, a single administration of miR-21 mimic enhanced the migration of pathogenic Th17 cells into the CNS by day 5 after transfer (Figures 6D and 6E). In contrast, no increase in transferred Th17 cell frequency was observed in the spleen or lymph nodes, although total cell numbers were reduced (Figure S6D). These findings indicate that elevated circulating miR-21 promotes the accumulation of pathogenic T cells within the CNS.

**Figure. 6:**
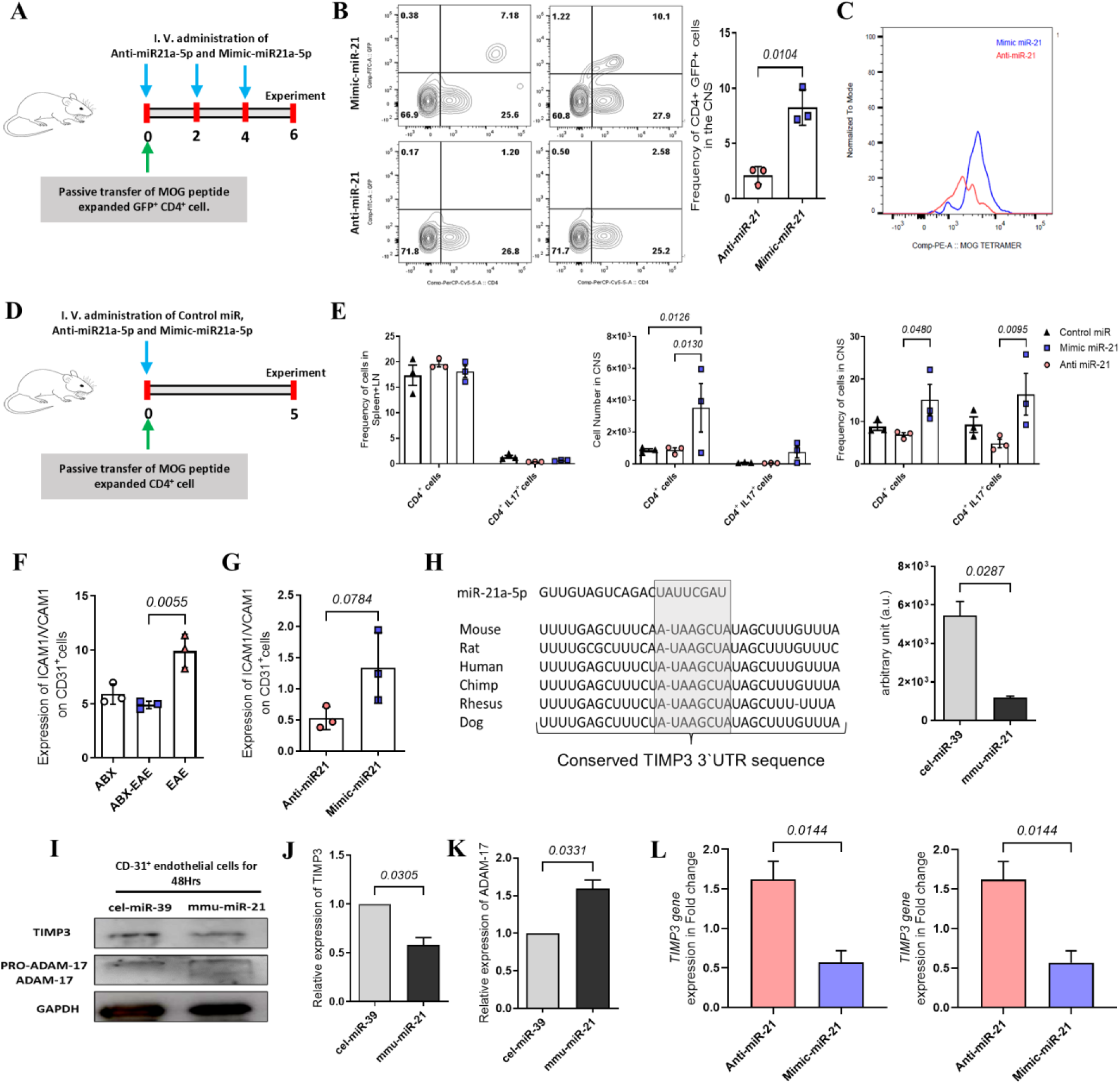
Cell-free miR-21 promotes the invasion of T cells into the CNS via targeting TIMP3. **A.** Experimental schematic. Mice receiving passive transfer of GFP-expressing CD4+ cells were administered miR-21 mimetic and anti-miR-21 oligonucleotides on days 0, 2, and 4. **B**. Promoted invasion of T cells to the CNS by increased circulating miR-21. Lymphocytes were isolated from the CNS of CD4^+^ GFP^+^ cell recipient mice and analysed for GFP+ cell frequency by flow cytometry. **C.** Representative of flow cytometer histogram showing the count of MOG tetramer^+^ CD4^+^ GFP^+^ cells in the CNS of recipient mice. (n = 3 independent experiments). **D.** Experimental schematic. Mice receiving passive transfer of CD4^+^ cells were administered Control miR, mimic-miR-21, and anti-oligo-miR-21 oligonucleotides on day 0. **E.** Frequency and cell count of CD4^+^ and IL-17^+^ cells in spleen+LN and CNS at day 5 post transfer (n = 3 independent experiments). **F.** Expression of cell adhesion molecules in CNS endothelial cells. Expression of ICAM1/VCAM1 on CD31^+^ CNS endothelial cells isolated on day 14 from EAE and ABX-EAE mice. **G.** Expression of cell adhesion molecules in CNS endothelial cells. Expression of ICAM1/VCAM1 on CD31^+^ CNS endothelial cells isolated from mice treated with miR-21-mimic and anti-miR-21 oligonucleotides on day 0, 2, and 4. (n = 3 independent experiments). **H.** Target gene for miR-21. The miR database (miRBase) indicated a predicted binding site for miR-21 in the 3΄-untranslated region (UTR) of TIMP3. Luciferase reporter gene assay showed that the reporter gene carrying the 3΄-UTR of TIMP3 was significantly suppressed in the presence of synthetic miR-21 (n=2 independent experiments). **I, J,** and **K.** Target protein expression CD31^+^ endothelial cells expressing TIMP3 and ADAM17 were transfected with miR-21 mimic and cel-miR-39 mimic (control) and cultured for 96 h. Cell lysate was prepared and examined by western blot analysis with antibodies against TIMP3, ADAM17, and GAPDH. **L.** qRT-PCR gene expression analysis. The expression of *TIMP3* and *ADAM17* genes in endothelial cells transfected with miR-21 mimic and anti-miR-21 oligonucleotides was examined by qRT-PCR (n = 3 independent determinations). Error bars represent the mean ± SEM values. In Figures B, G, J, K, and L, a two-tailed Mann-Whitney U test was used. In figures E and F, a 2-way ANOVA was used to calculate the *P* values.

Because endothelial adhesion molecules regulate leukocyte transmigration across the blood–brain barrier (BBB), we next examined ICAM1 and VCAM1 expression on CNS endothelial cells. Both molecules were significantly elevated in EAE mice compared with ABX-EAE mice (Figures 6F and S6E). Consistent with these observations, administration of miR-21 mimics increased endothelial ICAM1 and VCAM1 expression in the passive transfer model (Figure 6G), suggesting that circulating miR-21 promotes endothelial activation.

To identify downstream mediators of this response, we investigated ADAM17, a metalloprotease implicated in endothelial activation and leukocyte transendothelial migration [47, 48]. Protein interaction and pathway enrichment analyses linked ADAM17 to TNF signaling and leukocyte transendothelial migration pathways (Figures S6F-H). Screening of miRNA target databases identified a conserved miR-21 binding site within the 3′ untranslated region (UTR) of TIMP3, a physiological inhibitor of ADAM family metalloproteases (Figure 6H). Luciferase reporter assays confirmed direct interaction between miR-21 and the TIMP3 3′UTR (Figure 6H). In endothelial cells, miR-21 overexpression reduced TIMP3 expression and increased ADAM17 activation, whereas anti-miR-21 treatment had the opposite effect (Figures 6I-L). Protein interaction analysis further supported a regulatory relationship between TIMP3 and ADAM17 (Figures S4I and S6J). Collectively, these findings identify a miR-21–TIMP3–ADAM17 signaling axis that enhances endothelial activation and promotes the entry of pathogenic T cells into the CNS during autoimmune neuroinflammation.

### Circulating miR-21 is elevated in patients with multiple sclerosis

To determine whether the miRNA alterations identified in EAE were also present in human disease, we analyzed plasma samples from patients with multiple sclerosis (MS) and healthy controls (HC). Among the candidate miRNAs identified in EAE, only miR-21 was significantly elevated in MS patients compared with healthy controls, whereas miR-146a, miR-122, and miR-223 showed no significant differences between groups (Figure 7A). Because increased miR-21 expression in EAE was associated with *A. muciniphila* abundance and TLR2/TLR4 signaling, we next examined miR-21 expression in clinically defined MS subtypes. Plasma miR-21 levels were significantly increased in both relapsing-remitting MS (RRMS) and secondary progressive MS (SPMS) patients compared with healthy controls (Figure 7B). In contrast, serum TLR2/TLR4 agonist activity was reduced in MS patients relative to healthy controls (Figure 7C).

**Figure. 7:**
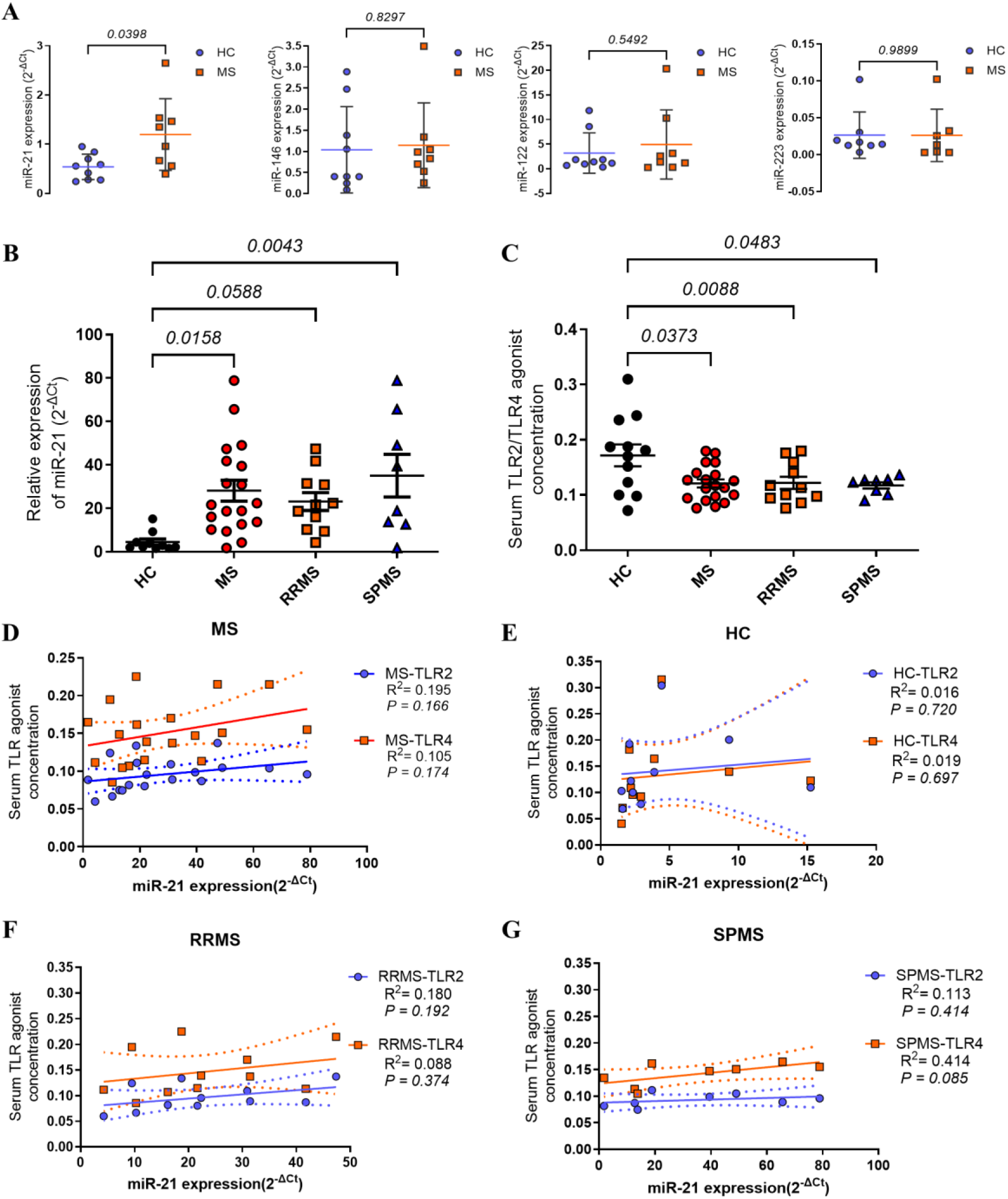
Blood circulating miR-21 in MS patients. **A.** Quantitative analyses of serum miRs. qRT-PCR analyses of the indicated miRNA in serum samples from patients with MS and healthy controls (HC) were performed (n=8 per group). **B.** Relative expression of serum miR-21. qRT-PCR analysis of miR-21 in serum samples of patients with MS (RRMS+SPMS), RRMS, SPMS, and healthy controls (HC) was performed (n = MS-19, RRMS-11, SPMS-8, HC-12, individual data). **C.** Quantitative analysis of serum TLR2/TLR4 agonist. TLR2- and TLR4-reporter cells were stimulated with an equal volume of MS patient serum. After 20 h of culture, the level of secreted embryonic alkaline phosphate (SEAP) was measured at 620 nm (n = 3 independent determinations). **D and G.** Correlation analysis of serum miR-21 and TLR agonist. Simple linear regression analysis was performed on MS patients’ serum concentrations of TLR2 and TLR4 agonists (Y axis) versus miR-21 expression (X axis). Error bars represent the mean ± SEM values. A Unpaired two-tailed Mann-Whitney U test, B,C, one-way ANOVA followed by Dunnett’s multiple comparisons test was used to calculate the *P* values.

To investigate the relationship between microbial-derived innate immune stimuli and miR-21 expression, we performed linear regression analyses comparing serum TLR2/TLR4 agonist activity with circulating miR-21 levels. Positive associations were observed between miR-21 expression and both TLR2- and TLR4-agonist activity in MS patients, whereas no such relationships were detected in healthy controls (Figures 7D and 7E). Similar positive trends were observed in both RRMS and SPMS subgroups, although these associations did not reach statistical significance (Figures 7F and 7G). Collectively, these findings demonstrate that circulating miR-21 is elevated in patients with MS and remains associated with serum TLR agonist activity, supporting the clinical relevance of the microbiota-miR-21 axis identified in EAE.

## Discussion

Research on non-coding RNAs has established miRNAs as central post-transcriptional regulators of immune and inflammatory gene networks. Multiple profiling studies have reported altered miRNA signatures in MS; however, the upstream regulators and the mechanisms underlying disease amplification remain incompletely defined. Building on prior observations that miR-21 promotes Th17 differentiation and that microbiota composition influences EAE, our study integrates these fields by identifying a microbiota-driven miR-21 pathway that links gut dysbiosis to BBB dysfunction and CNS autoimmunity Our study integrates these concepts by identifying a microbiota-driven miR-21 pathway that mechanistically links gut dysbiosis to BBB dysfunction and CNS autoimmunity. We demonstrate that *Akkermansia muciniphila*–derived lipopolysaccharides activate intestinal TLR2/4 signaling, inducing epithelial miR-21 secretion that enhances Th17 polarization and compromises BBB integrity through the TIMP3–ADAM17 axis. We and others have reported altered miRNA expression profiles in MS, yet the results are notably inconsistent across studies [6, 7, 11]. Such variability may arise from differences in disease stage and environmental factors, particularly variations in gut microbiome composition. Because microbiota can modulate host miRNA expression, we investigated how microbial dysbiosis influences autoimmune demyelination. Antibiotic-induced microbiome depletion suppressed EAE and significantly reduced the expression of miR-21, miR-146a, and miR-223. Notably, among these candidates, only inhibition of miR-21 substantially attenuated disease severity and decreased CNS T-cell infiltration, confirming miR-21 as a key pathogenic miRNA in EAE. These findings refine earlier descriptive studies by functionally identifying miR-21 as a dominant pathogenic driver rather than a secondary inflammatory marker.

Previous work demonstrated that germline deletion of miR-21 confers resistance to EAE by impairing Th17 differentiation [9]. In line with these findings, we show that miR-21 enhances IL-17 and GM-CSF production under Th17-polarizing conditions. However, our data extend prior knowledge by demonstrating that circulating miR-21 induction during EAE is microbiota-dependent and not solely derived from adaptive immune cells. To determine this, we examined germ-free mice. Germ-free mice failed to upregulate miR-21 following immunization, indicating that microbial factors control its expression. Research indicates that in vitro CD4+ T cells express miR-21, and that germ-free mice harbor a decreased CD4+ cell population. Therefore, to explore additional in vivo sources of miR-21, we utilized the Rag1-deficient model, which lacks mature T and B cells. Remarkably, miR-21 remained inducible following immunization in Rag1-deficient animals, suggesting that microbial signals drive its induction in early EAE.

Commensal bacteria are known to trigger epithelial miR-21 during barrier stress, prompting us to assess intestinal sources in the EAE setup. We found a marked induction of miR-21 in intestinal epithelial cells by day 7 of EAE, coinciding with increased local CD4^+^ T-cell infiltration and elevated expression of IL-17 and GM-CSF, key cytokines in EAE pathogenesis [49]. Prior studies have shown that encephalitogenic T cells transiently reside in the intestine before CNS entry, and that restricting gut migration reduces EAE severity. Our findings provide a mechanistic framework that integrates these observations by identifying epithelial-derived miR-21 as a potential molecular mediator linking early gut immune activation to subsequent CNS pathology.

Because miRNAs circulate in stable complexes or exosomes [50]. Previous studies have shown that exosomal miRNAs can modulate endothelial function in inflammatory diseases [51]. Considering this concept, we examined whether exosomal miR-21 directly contributes to EAE progression. EAE mice exhibited a pronounced increase in miR-21-enriched exosomes, which was attenuated by antibiotic treatment. Fluorescently labeled exosomes transferred from EAE to control mice preferentially accumulated in CNS endothelial layers, suggesting a potential route for systemic miRNA-mediated barrier disruption. A MOG peptide-primed CD4+ T cell passive transfer experimental model confirmed this: intravenous administration of miR-21 mimics enhanced, while antisense inhibition reduced, the migration of pathogenic Th17 T cells into the CNS, which is responsible for EAE induction. Inflammatory cytokines are known to promote CNS infiltration by up-regulating endothelial adhesion molecules ICAM-1 and VCAM-1 [52]. Consistent with this finding, we observed increased ICAM-1 and VCAM-1 expression in the CNS of EAE and miR-21 mimic–treated mice, whereas antibiotic or anti-miR-21 therapy suppressed their expression. Collectively, these observations position miR-21 as an active regulator of endothelial activation and immune-cell trafficking across the BBB. So, exosomal miR-21 influences endothelial integrin to support T-cell invasion; however, the target genes of miR-21 are unknown.

In recent years, ADAM metalloproteinases have attracted significant attention for their roles in immunity, particularly ADAM17, a TNF-converting enzyme in neuroinflammatory conditions, including MS [48]. Therefore, given ADAM17’s role in MS, to identify the molecular mechanisms underlying this effect, we focused on ADAM17. We identified TIMP3, a well-characterized endogenous inhibitor of ADAM17, as a potential miR-21 target. The nucleotide sequence of mature miR-21 complements the 8-mer conserved 3′-UTR binding sites of the transcriptional inhibitor metal protease enzyme TIMP3. ADAM17 is a membrane-anchored shedder that cleaves multiple substrates, including TNF, TNF receptors, and adhesion molecules such as ICAM-1 and VCAM-1. These cleavage events are known to increase vascular permeability and promote leukocyte adhesion and migration across the BBB called transendothelial migration [53, 54]. Therefore, we speculated that exosomal miR-21 deposited in endothelial cells may modulate the biological function of TIMP3, a key inhibitor of several proteases, particularly ADAM17; we experimentally confirmed the miR-21 target-binding site on the TIMP3 mRNA. However, initially, to understand their co-existence, we performed database analysis. Here, STRING and KEGG databases confirmed the association of TIMP3, ADAM17, and TNF protein expression; therefore, to establish this association among proteins regulated by miR-21, we transfected endothelial cells with miR-21 mimics and performed western blotting. Overexpression of miR-21 markedly reduced TIMP3 protein levels while increasing the abundance and activation of pro-ADAM17, consistent with proteolytic disinhibition. Conversely, inhibition of miR-21 by anti-miR-21 oligonucleotides restored TIMP3 and stabilized endothelial junctions. Importantly, exposure of endothelial cells to exosomal miR-21 isolated from EAE serum reproduced this effect, confirming that circulating miR-21 can be functionally internalized by the BBB endothelium. Hyperactive ADAM17 subsequently enhances TNF and ICAM-1 shedding [55, 56], amplifying cytokine signaling and further compromising tight-junction integrity. Together, these findings establish a mechanistic cascade in which microbiota-induced miR-21 suppresses TIMP3, leading to unchecked ADAM17 activation, the excessive release of soluble TNF and adhesion molecules, and progressive destabilization of the BBB. Thus, miR-21 operates at a critical junction between immune activation and vascular dysfunction, amplifying peripheral inflammation and facilitating immune cell infiltration into the CNS.

In the prodromal phase of EAE, the compromised gut barrier and elevated epithelial miR-21 expression suggested a direct microbiome-driven inflammatory signal. This aligns with evidence that miR-21 knockout mice resist DSS-induced colitis [57–59], reinforcing the intestinal origin of its proinflammatory effects. The gut microbiota undergoes marked changes during EAE [60], and our data revealed a transient yet pronounced expansion of the mucin-degrading bacterium *A. muciniphila* early after immunization (day 7), coinciding with miR-21 induction. In the antibiotic-treated EAE mice, *L. reuteri* was abundant, whereas *A. muciniphila* abundance was reduced, and the microbial diversity recovered at day 14 of EAE. Further deep sequencing helped us to identify the strain resemblance to the previously identified *A. muciniphila* JCM 30893 in a healthy Japanese cohort [61]. Therefore, we challenged the role of the *A. muciniphila* JCM 30893 and isolated *L. reuteri* on EAE induction. Colonization of ABX-treated SPF mice with the typed *A. muciniphila* JCM 30893 strain exacerbated EAE symptoms and elevated IL-17 expression in CNS CD4+ T cells. We also observed a significant increase in circulatory miR-21 in both EAE and colonized ABX-EAE mice. On the other hand, mono-colonizing GF mice with *A. muciniphila* JCM 30893 and isolated L. reuteri do not raise serum miR-21 levels in mice, clearly indicating that miR-21 generation is indirectly immunization-dependent. This result, implicating *A. muciniphila* as the inducer, suggests it exacerbates Th17-driven pathology through miR-21.

This observation resonates with clinical studies reporting increased fecal abundance of *A. muciniphila* in MS patients[62], but its role in inducing inflammation remains controversial. Transferring microbiota from MS patients enriched in *A. muciniphila* aggravates EAE in mice, and in an in vitro setup, it enhances IL-17 production in human PBMCs [21, 27]. In contrast, a study on fecal miRNAs in EAE demonstrated that increased fecal miR-30d levels were associated with increased *A. muciniphila* and promoted Treg differentiation, thereby ameliorating EAE [10, 30]. Such divergent findings likely reflect strain-specific or context-dependent bacterial phenotypes, supported by recent evidence that bacterial genomes can undergo reversible rearrangements affecting virulence and host interactions [63]. Our findings suggest that under inflammatory conditions, *A. muciniphila* acquires a proinflammatory phenotype capable of inducing miR-21 expression or gut mucosal barrier disruption is key for such pathology.

Metagenomic functional analysis revealed upregulation of bacterial secretion and LPS biosynthesis genes in EAE, consistent with increased abundance of *A. muciniphila* in early EAE. Prior work demonstrated that though *A. muciniphila* is gram-negative, still it can activate TLR4 and TLR2 receptors and its phospholipids can induce immunomodulatory responses through TLR2–TLR1 heterodimers [64, 65]. We confirmed the atypical characteristics of LPS using TLR2- and TLR4-expressing reporter cell lines with both heat-inactivated and purchased *A. muciniphila* LPS. Since we observed compromised gut permeability and increased *A. muciniphila,* along with upregulation of the LPS biosynthesis pathway, we hypothesized that its atypical LPS may activate epithelial TLR2 and TLR4, thereby inducing miR-21. To test this, we used an in vitro system with the murine epithelial cell line STC-1; indeed, heat-inactivated *A. muciniphila* and its purchased LPS induced robust miR-21 secretion compared with *LPS from L. reuteri and E. coli*, respectively. Further, blocking either TLR2 or TLR4, or both using inhibitors, in the presence of *A. muciniphila* LPS significantly reduced miR-21 release, confirming that co-stimulation of TLR2 and TLR4 is required for the amplified secretion of miR-21. In GF mice, mono-colonization with *A. muciniphila* does not induce miR-21; by contrast, EAE-induced GF mice were administered purified *A. muciniphila* LPS, resulting in a dramatic upregulation of serum miR-21. This shows that disruption of the gut barrier may be necessary for LPS-induced miR-21 generation in vivo. Finally, to track the exosome migration and inflammatory property, the exosomes generated through *A. muciniphila* LPS stimulation on STC-1 cell lines were transferred to SPF mice, resulting in preferential deposition in CNS tissues but not the CpG-stimulated exosomes. In vitro setup: adding LPS-derived exosomes to CD4+ cells increases IL17 and GMCFS secretion. These results highlight a possible route by which microbial signals can reach the brain and regulate inflammation during EAE [62]. Collectively, these data support the notion that *A. muciniphila*–derived LPS triggers TLR2/TLR4-dependent secretion of miR-21, linking microbial dysbiosis to neuroinflammatory signaling.

Although MOG-induced EAE is an imperfect MS model for microbiome analysis, it recapitulates core inflammatory mechanisms relevant to disease progression [66, 67]. Several researchers have reported the involvement of diverse miRNAs in MS and EAE; however, it is necessary to identify shared miRNAs across these conditions to understand their roles in autoimmunity. Therefore, we initially selected serum samples from untreated or DMT-treated MS patients to eliminate drug effects on miRNA expression and to screen for all upregulated EAE miRNAs. Consistent with our murine data, serum miR-21 levels were significantly elevated in both untreated and DMT-treated MS patients, and its expression correlated positively with disease severity across both relapsing-remitting MS (RRMS) and secondary-progression MS (SPMS) subtypes. Our prior microbiome analysis identified higher *A. muciniphila* abundance in RRMS, suggesting a potential shared microbe-miRNA axis in human MS. Interestingly, while serum TLR4 agonist levels were reduced in MS patients, this likely reflected receptor activation and ligand consumption. However, in MS and its subtypes, the correlation between TLR4 and TLR2 ligand levels and serum miR-21 expression was positive, whereas in SPMS patients, the correlation was nearly statistically significant, indicating ongoing immune engagement with microbial components. These findings suggest that increased *A. muciniphila* in MS may sustain epithelial and systemic TLR signaling, thereby maintaining elevated miR-21 levels even as ligand concentrations decline.

Taken together, our results delineate a microbe-miRNA-endothelium axis that links gut dysbiosis to CNS inflammation. We propose that *A. muciniphila*–derived LPS co-activates TLR2 and TLR4 in intestinal epithelial cells, driving miR-21 production. Circulating and exosomal miR-21 amplify Th17 polarization and disrupt BBB integrity via the TIMP3–ADAM17 pathway, facilitating Th17 cell migration into the CNS. This integrated mechanism reconciles conflicting evidence about the dual nature of *A. muciniphila* in MS and positions miR-21 as a central effector of the gut–brain axis, offering a promising RNA-targeted therapeutic strategy for autoimmune neuroinflammation.

## Limitations of the study

Our study has certain limitations. We propose that LPS from *A. muciniphila* coactivates TLR2 and TLR4 to induce epithelial miR-21 secretion, but it remains unclear whether other gut bacteria with atypical LPS structures or microbial components exert similar effects. Future research should employ strain-level profiling and functional screening to identify bacterial taxa capable of modulating host miRNA expression. As global miR-21 knockout confers complete EAE resistance, it likely affects multiple targets beyond those identified here; thus, its influence on non-immune and stromal cell signaling requires further study. Finally, our data in SPF mice suggest that miR-21 may regulate additional immune pathways that amplify T cell–mediated autoimmunity, warranting deeper mechanistic exploration.

## Funding

MM is grateful for support from the Multiple Sclerosis International Federation (MSIF) through the McDonald Fellowship, sponsored by ARSEP and the Japanese Multiple Sclerosis Society (JMSS). The authors thank Dr. Benjamin Raveney for helpful advice and support with the experimental design and protocols. The authors thank H. Yamaguchi, A. Takeo, K. Inoue, C. Koto for technical support. This work was supported by a practical research project for rare/intractable diseases from the Japan Agency for Medical Research and Development (AMED) (Grant #: 17ek0109190h0002; 20ek0109315h0003), AMED-Core Research for Evolutional Science and Technology (CREST) (Grant #:18gm1010011h0001; 19gm1010011 h0002), and AMED-CREST (Grant Number JP21gm1010006) to H.M., J.K., and A.T.

## Author contributions

TY, SO, and MM conceptualized the study; MM conceived this study, designed and performed the experiments, analyzed the data, and wrote the manuscript. DT isolated the bacteria and helped with data analysis. HH supervised and assisted with miRNA isolation, quantification, and other miRNA-related experiments. WS collected samples from both MS and healthy individuals. HM, KH, JK, and AT sequenced and analyzed the gut microbiome data. TY, WS, and SO supervised the experiments; TY, HH, and KK revised and finalized the manuscript. All the authors have read and approved the final manuscript.

## Competing interests

The authors declare no competing interests.

## Materials & Correspondence

Further information and requests for resources and reagents should be directed to and will be fulfilled by the lead contact, **Takashi Yamamura**.

---

**Figure. S1:**
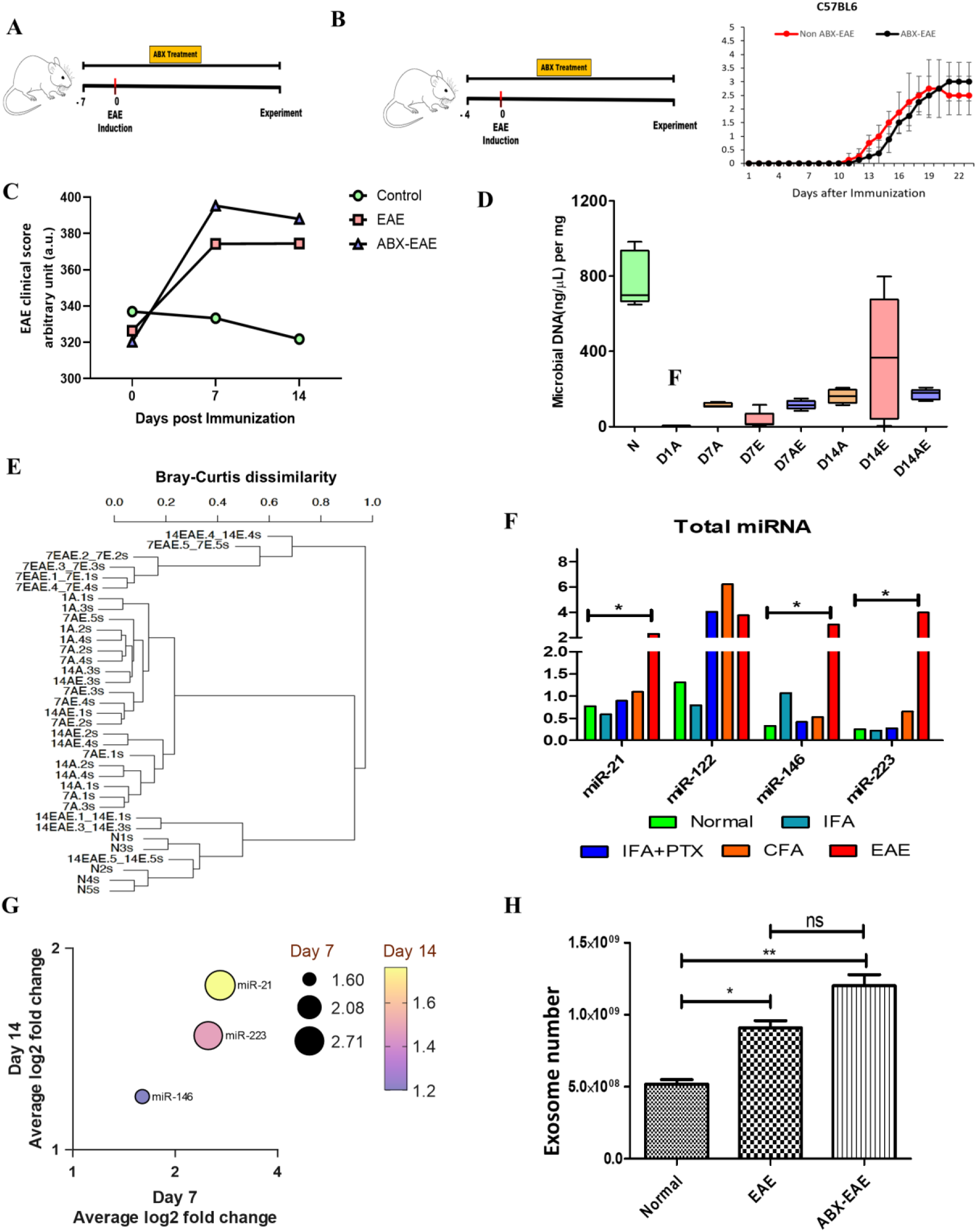
Change in gut bacteria affects EAE severity. **A**. Schematic representation. Generation of gut microbiome dysbiosis model mice (ABX-EAE), Mice were orally administered an antibiotic cocktail (ABX). Every day, starting from one week before immunization and continuing till the date of the experiment. **B.** Schematic representation. Mice treated for 4 days pre-immunization do not change the EAE clinical score. **C.** Gut barrier dysfunction. Leaky gut analysis at day 7 and 14 postimmunization in Control, EAE, and ABX-EAE mice was performed by orally administering 4 KDa FITC dextrin; after 4 h of administration, the serum sample was collected and fluorescence was measured at 530nm. (n=2 in control and n=3 in EAE and ABX-EAE), auto fluorescence was measured and normalized before FITC dextrin administration. **D.** Fecal DNA quantification. Fecal microbial DNA content in normal(N), dysbiosis (AE) and intact(E) mice, significant reduction in the microbial DNA content in dysbiosis and intact day 7 mice, but this was recovered in day14 intact mice but this was recovered in day 14 intact mice. **E.** Bray–Curtis dissimilarity showed fecal microbial dissimilarity among normal (N), EAE (E), ABX-EAE (AE), and ABX-treated non-EAE (A) models. **F.** qRT-PCR miRNA quantification. Expression of four known miRNAs, namely miR-21, miR-122, miR-146 b, and miR-223, was analysed in different combinations of EAE induction components. upregulation of liver-associated miRNA miR-122 in CFA without peptide treatment, but no other miRNA expression was increased. **G.** Multiple variable analysis. Graph representing average log 2-fold change miR-21 expression at day 7 and day 14 EAE. (n=3 independent mice). **H.** Exosome quantification. Circulating serum exosome numbers were quantified in the normal, EAE, and antibiotic-treated EAE mice. There is a significant increase in exosomes observed in EAE and ABX-EAE compared with normal mice. Error bars represent the mean ± SEM values; an a two-tailed Mann-Whitney U test was used to calculate the *P* value.

**Figure. S2:**
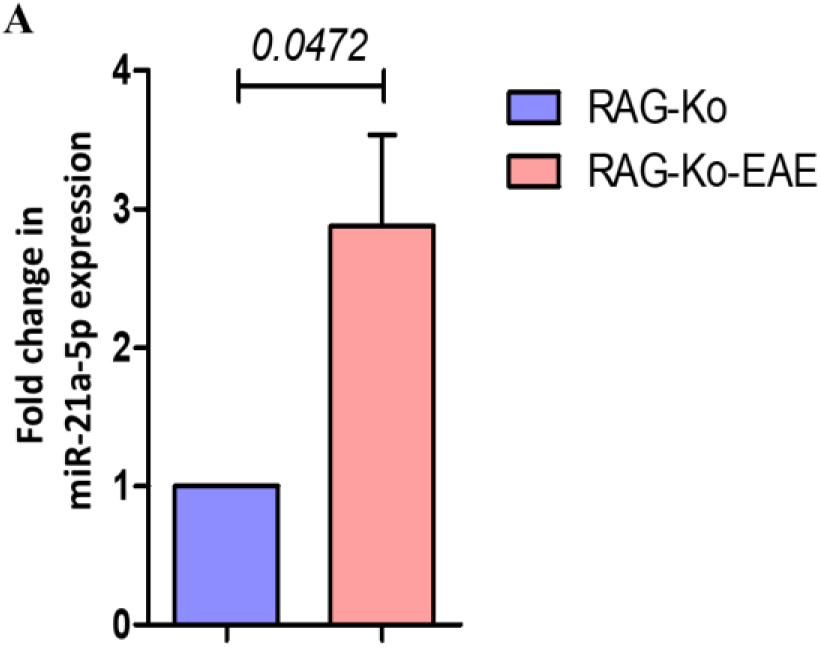
Microbiome-dependent miR-21 expression. **A.** qRT-PCR miRNA quantification. Serum levels of circulating miR-21 in MOG-immunized RAG-Ko mice at day 18. Error bars represent the mean ± SD values. An a two-tailed Mann-Whitney U test was used to calculate the *P* value.

**Figure S3:**
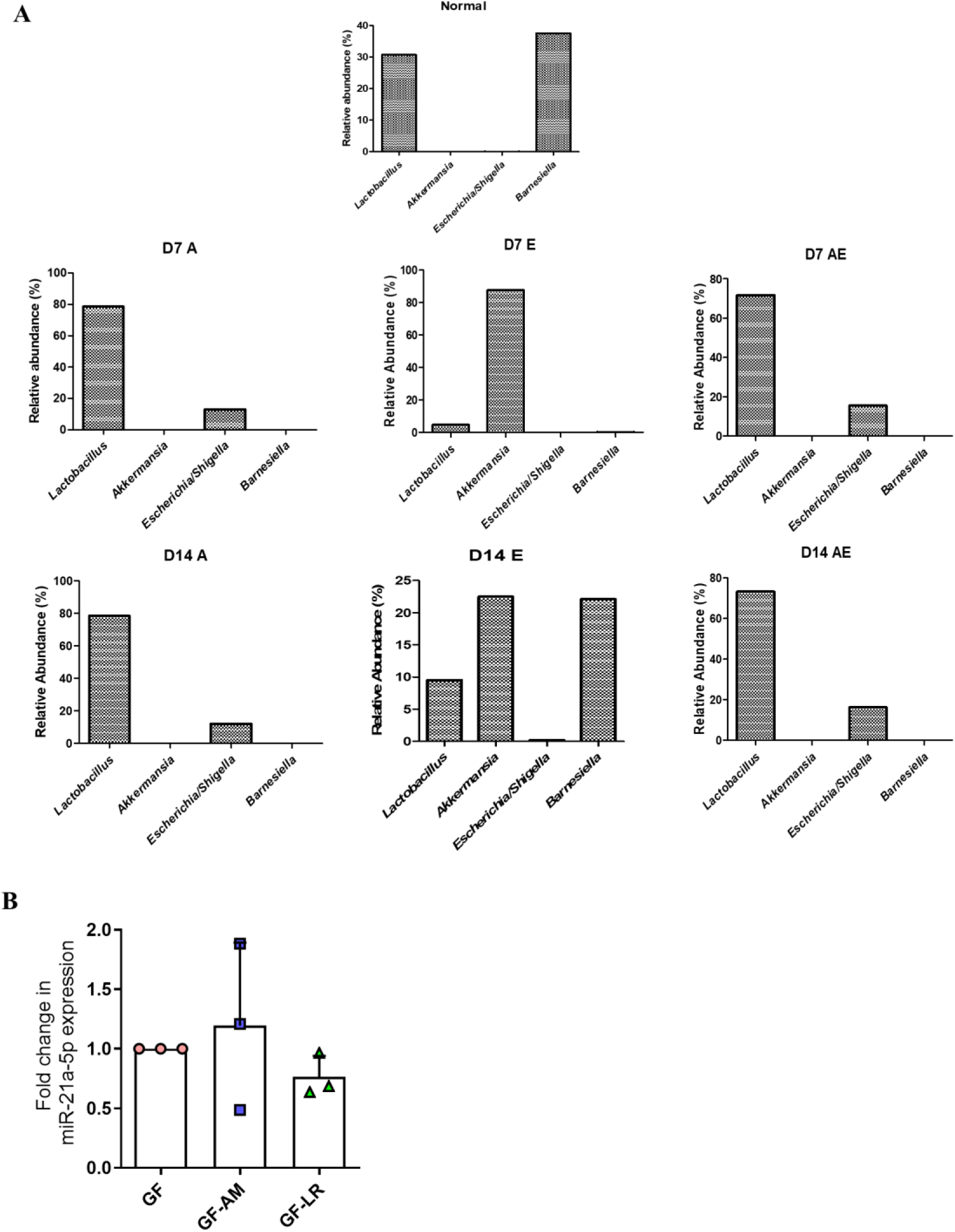
differential microbial population. **A.** Relative abundance of bacterial species. Four highly expressed gut bacteria in Normal, Antibiotic administration(A), Antibiotic administered EAE(AE), and PBS-treated EAE(E) mice at day 7 and day 14 from the date of immunization. **B.** qRT-PCR analysis of blood circulating miR- 21. Freshly collected serum from Control (GF) *A. muciniphila* monocolonized GF mice (GF-AM) and *L. reuteri* monocolonized GF mice (GF-LR) was used to quantify circulating miR-21. (n=3 independent experiments). Error bars represent mean ± SEM values. An ordinary 2-way ANOVA was used to calculate the *P* values.

**Figure S4:**
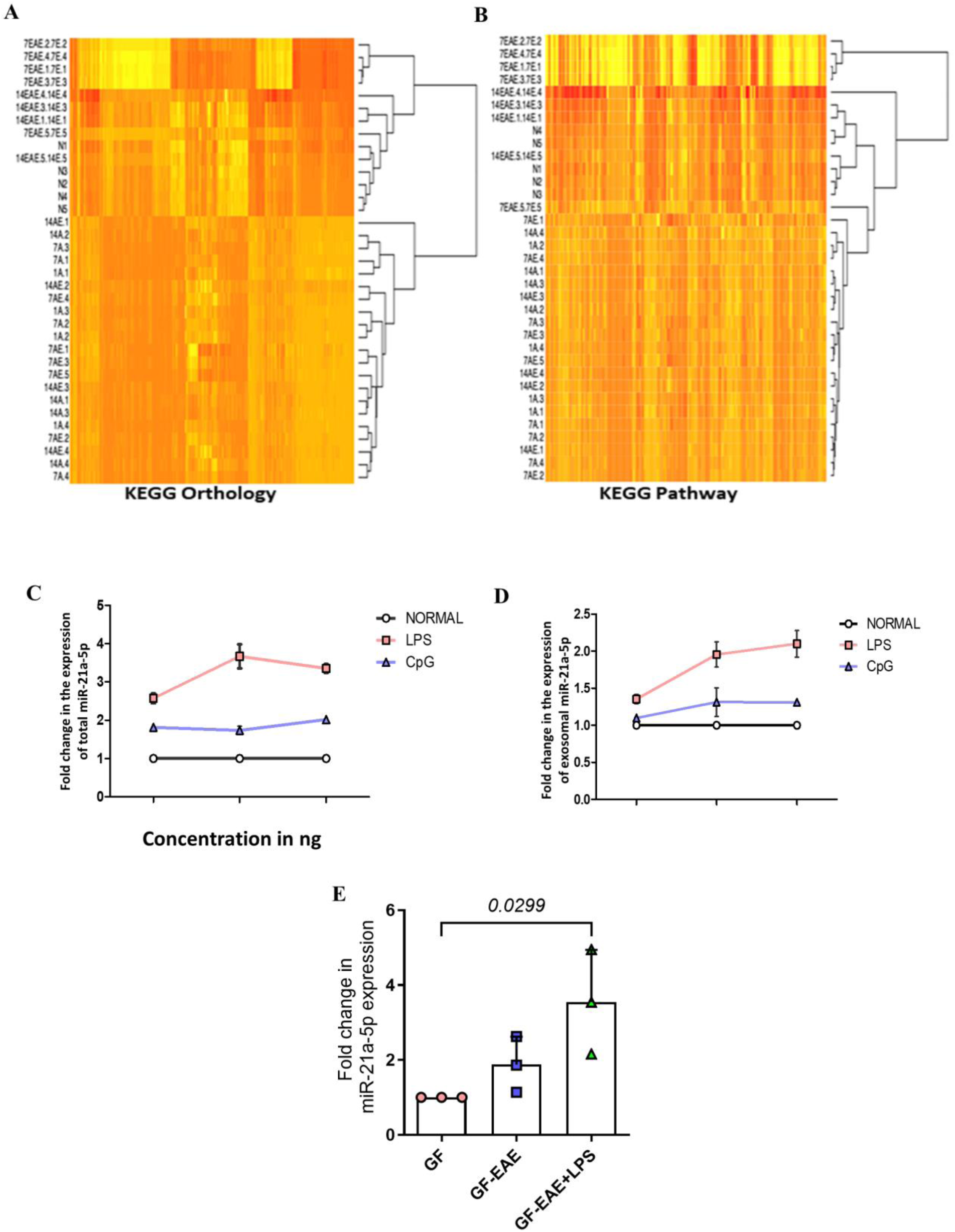
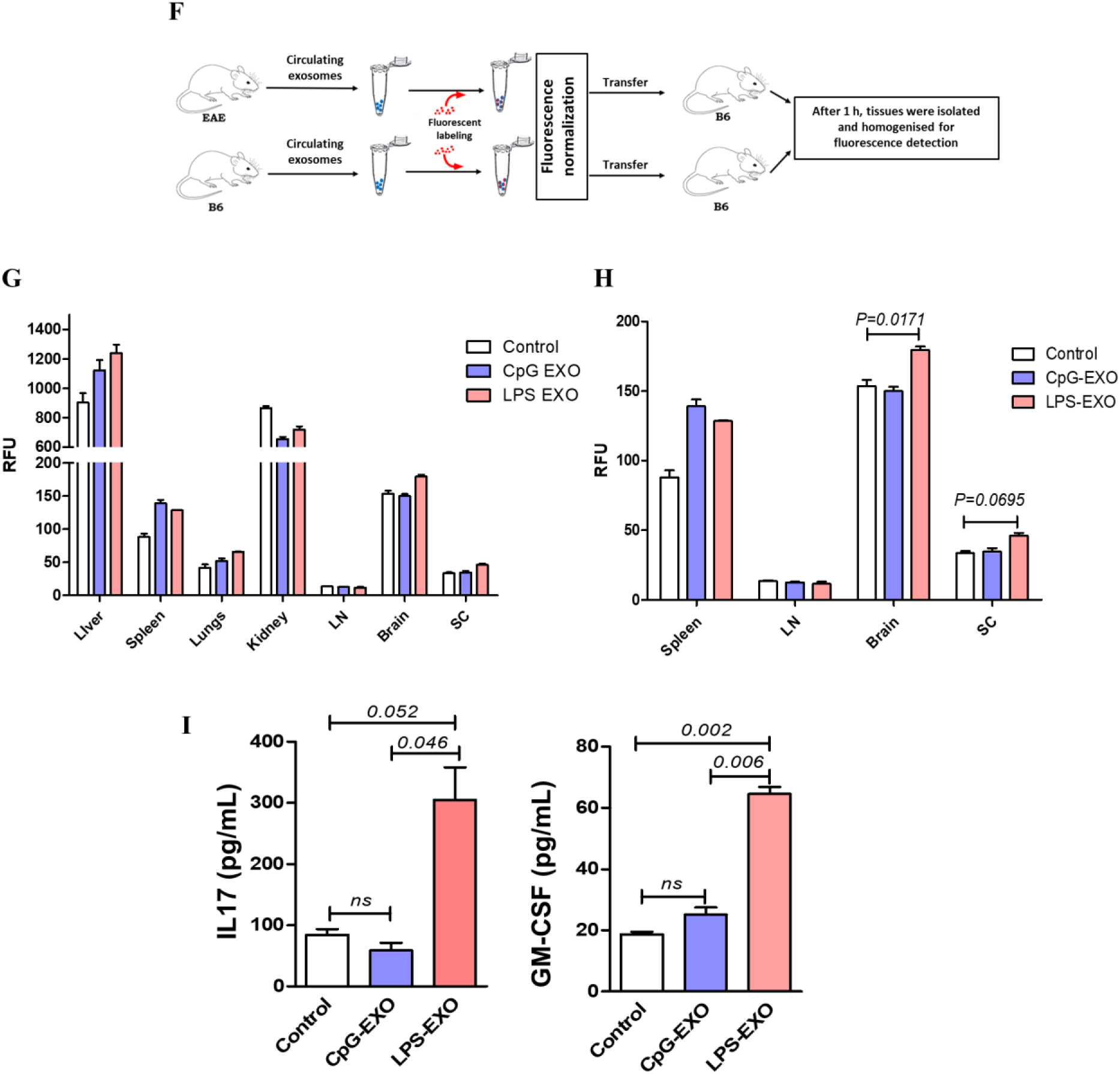
*A. muciniphila* LPS induces cell-free and exosomal miR-21 through TLR. **A-B.** Difference between fecal microbial KOs. Heat map representing KEGG orthology and KEGG pathway enrichment analysis between Normal, Antibiotic administration(A), Antibiotic administered EAE(AE), and PBS-treated EAE(E) mice at day 7 and day 14 from the date of immunization. **C-D**. qRT-PCR miRNA quantification. Stimulation of TLR2 and TLR4 of the STC-1 epithelial cell line with atypical LPS showed increased cell-free and exosomal miR-21. **E.** qRT-PCR analysis of blood circulating miR-21. Freshly collected serum from experimental mice was Control (GF), EAE induced GF mice (GF-EAE) and EAE induced and LPS administrated GF mice (GF-EAE+LPS) used to quantify circulating miR-21 on day 10 post-immunization. (n=3 independent experiments). **F-H**. Schematic representation and exosome tracking analysis. Exosomes derived from normal epithelial cells and LPS, CpG-stimulated epithelial cells were isolated and labelled with fluorescence dye and transferred to normal SPF mice. We harvested different tissue samples after 1 hour of transfer and read the fluorescence. We found significantly increased exosome deposition in many tissues, including the brain and spinal cord. **I.** Intracellular cytokine analysis. Naive CD4^+^ cells were cocultured with exosomes derived from lipopolysaccharide (LPS)- and CpG-stimulated epithelial cells for 48 h in a Th17-conditioned medium. Cytokines released into the medium were analyzed using ELISA. (n = 3 independent determinations). Results and data represent 2 independent experiments. Error bars represent the mean ± SEM values; in figures E, G-I, a 2-way ANOVA was used to calculate the *P* values.

**Figure. S5:**
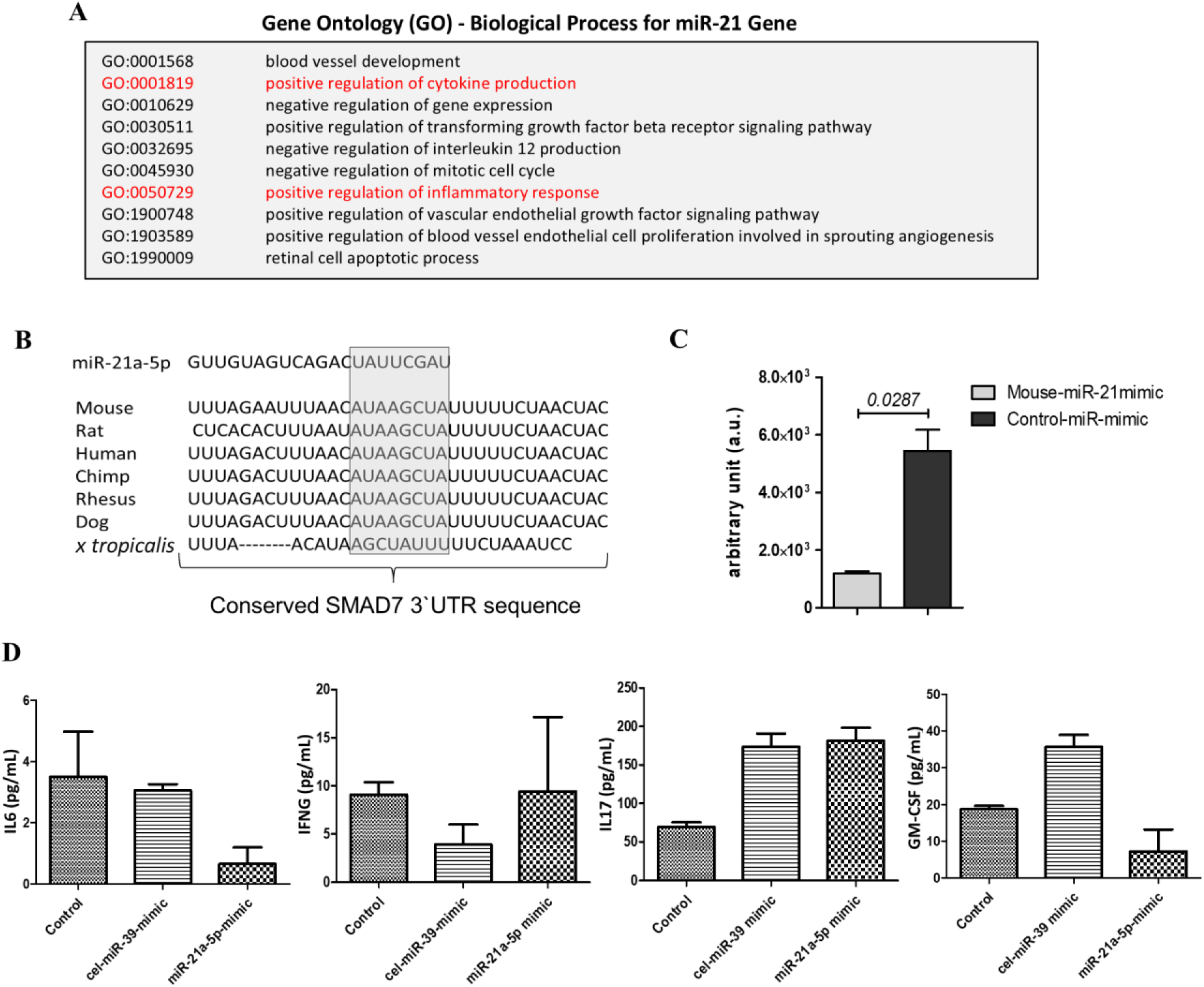
miR-21 influences T cell pathogenicity in EAE. **A.** Gene ontology analysis. It revealed that miR-21 is a positive regulator of cytokine production and the inflammatory response. **B.** miRNA target database. miR database (miRBase) shows the highly conserved binding site of miR-21 in the 3′-untranslated region (UTR) of SMAD7. **C.** Luciferase assay for SMAD7. Suppression of the 3′-UTR of the SMAD7 gene (mentioned in F) present in Goclone® reporter vectors with synthetic miR-21. Results of luciferase assay confirming the binding site of miR-21 in SMAD7 using reporter gene luciferase expression with its control (n=2 independent experiments). **D.** Intracellular cytokine analysis using ELISA. Naïve CD4+ T cells of normal B6 mice were transfected with miR-21 mimics and cultured in Th1 conditional media. Here, no significant differences in cytokines INFγ, IL-6, IL-17, and GM-CSF were observed in Th1-conditioned medium. (n=3 independent determinations) Error bars represent the mean ± SD values.

**Figure S6:**
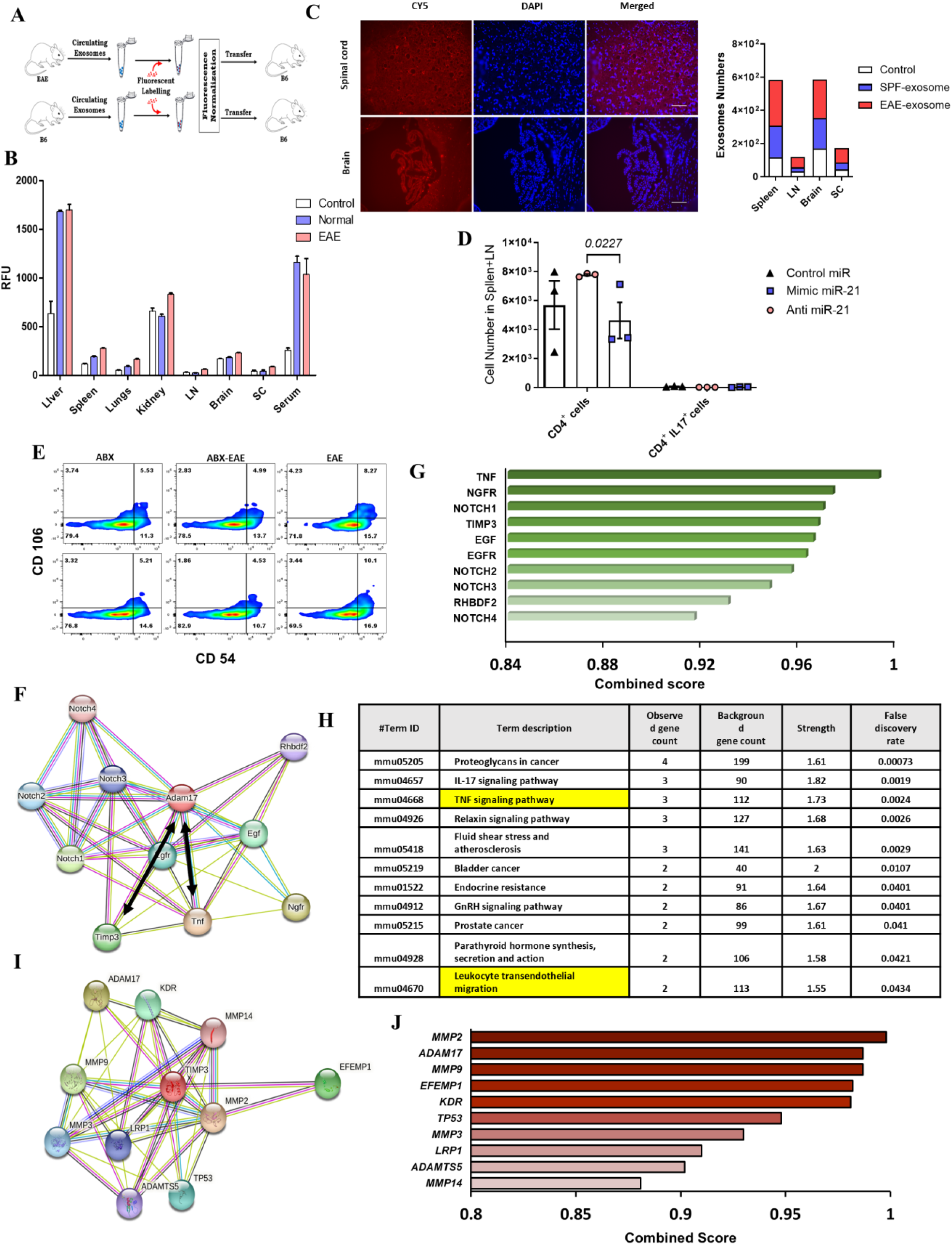
Circulating miR-21 promotes T cell invasion to CNS. **A.** Schematic representation. Exosomes from control and EAE mice (peak disease) were isolated and labeled with exosome fluorescent dye (ExoGlow™) and transferred to control B6 mice. After 1 h, tissues were harvested, homogenized, and analyzed for fluorescence. **B.** Exosomes quantification. Exosomes from EAE and Normal mice were isolated and labelled with fluorescence dye and transferred to normal SPF mice. We harvested tissue samples after 1 hour of transfer and measured fluorescence; we found significantly increased exosome deposition in many tissues. **C.** Histological analysis. Analysing the brain and spinal cord using histology, the deposition of labeled exosomes in the SC and brain of recipient B6 mice. (n=2 independent experiments, scale bar represents 100 μm). Differences in fluorescence emission were observed in different tissues, including the brain and spinal cord. Exosomes from EAE and normal mice were isolated and labelled with fluorescence dye and transferred to normal SPF mice. We harvested tissue samples after 1 hour of transfer and measured fluorescence; we found significantly increased exosome deposition in many tissues. **D.** Cell count of CD4^+^ and IL-17^+^ cells in spleen+LN at day 5 post transfer (n = 3 independent experiments). **E.** Flow cytometer analysis. Expression of ICAM1/VCAM1 on CD31^+^ CNS endothelial cells isolated from day 14 of EAE, and ABX-EAE mice. **F and G.** String analysis for protein-protein interaction. This showed that the proteins associated with ADAM-17, TNF, and TIMP3 are significantly associated with ADAM-17 expression. **H**. KEGG pathway analysis. List of KEGG pathways associated with ADAM-17, the TNF signalling pathway, and leukocyte transendothelial migration are strongly associated with ADAM-17 expression. **I and J**. String analysis for protein-protein interaction. This shows the correlation in the expression of the TIMP3 and ADAM17 genes, with a minimum interaction score of 0.700 (high confidence).In Figure D, a 2-way ANOVA was used to calculate the *P* values.

**Figure S7:**
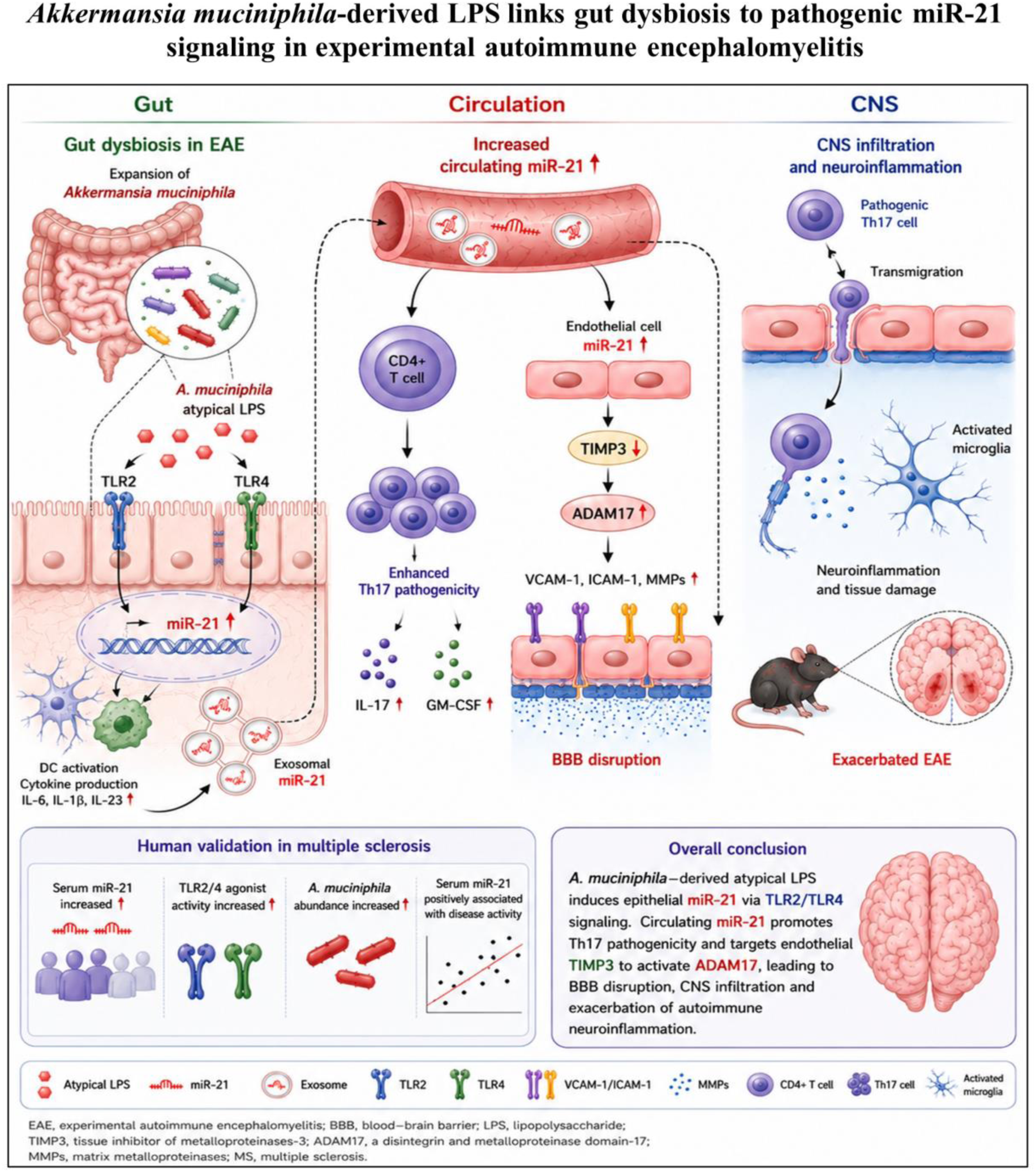
Molecular mechanisms underlying the roles of *Akkermansia muciniphila* and miR-21 in experimental autoimmune encephalomyelitis.

**Table S1:** Demography of MS patients and Healthy serum samples used for miRNA expression analysis.

| **Diagnosis** | **MS** | **HC** |
| --- | --- | --- |
| Number of Samples | 8 | 8 |
| Date of sample collection | < 2 years | < 4 years |
| Average Age | 48± 9 | 41.1± 5.8 |
| Sex | F-7<br>M-1 | F-6<br>M-2 |
| Diagnosis (more latest ) | RRMS-6<br>SPMS-1<br>PPMS-1 | - |
| Disease Duration | 6.9 years | - |
| EDSS (±SD ) | 4.12± 1.55 | - |
| ARR | 1.125± 0.9 | - |

**Table S2:** Demography of MS patients and Healthy serum samples used for miRNA expression analysis.

| Diagnosis | MS | HC |
| --- | --- | --- |
| Number of Samples | 19 | 12 |
| Date of sample collection | < 3 years | < 4 years |
| Average Age | 45±15 | 42.9 ± 5.8 |
| Sex | F-13<br>M-6 | F-9<br>M-3 |
| Diagnosis (more latest ) | RRMS-6<br>SPMS-1<br>PPMS-1 | - |
| Disease Duration | 6.9 years | - |
| EDSS (±SD ) | 2.92 ± 2.55 | - |
| ARR | 0.85 ± 1.2 | - |

## Notes

### Competing Interest Statement

The authors have declared no competing interest.

